# Dinucleotide codon mutations as a marker of diversifying selection in *M. tuberculosis*

**DOI:** 10.1101/2025.05.30.657033

**Authors:** Danila Zimenkov, Anastasia Ushtanit

**Affiliations:** Engelhardt Institute of Molecular Biology, Russian Academy of Sciences, 119991 Moscow, Russia

**Keywords:** tuberculosis, drug resistance, evolution, dinucleotide mutations, virulence, rifampicin, bedaquiline, linezolid

## Abstract

The evolution of the human pathogen *M. tuberculosis* is shaped by various but interconnected processes of drug treatment pressure and host adaptation. We hypothesize that rarely accounted dinucleotide substitutions within a single codon, which allow for a broader range of amino acid substitutions than single nucleotide changes, are a significant aspect of diversifying selection.

From the analysis of 43 studies, comprising 11,730 clinical isolates with resistance to rifampicin, 11 different dinucleotide substitutions were identified in 54 codons of resistance-determining regions of the *rpoB* gene. The prevalence of such substitutions is approaching 4%. Although rifampicin was introduced in treatment regimens in the 1970s, dinucleotide substitutions were also found in resistance determinants for newer drugs, linezolid and bedaquiline, *rplC*, and *atpE*, despite the significantly smaller number of resistant clinical isolates reported.

Conducting a genome-wide analysis of dinucleotide mutations in the dataset of 9,941 genomes studied by the CRYpTIC Consortium, in addition to resistance determinants, we discovered three genes with a significantly elevated number of dinucleotide substitutions, which are presumably related to virulence and host adaptation. Two substitutions, *cyp138* P114F and L191A are supposed to occur early in the evolutionary history of lineage 2 and are now under strong selection for reverse substitutions. Two amino acid substitutions in the third gene, *rv2024c* N508T and C514L, could also be obtained by single nucleotide changes and therefore are supposedly being selected based on frequency of codon usage.

The signature of dinucleotide mutations introduces a novel approach to understanding the evolution of pathogen and identifying potential targets for antivirulence drugs. They underscore the complexity of the evolutionary dynamics within this pathogen, driven by diverse selection pressures, shedding light on the ongoing battle between *M. tuberculosis* and its human host.

## 1 Introduction

Tuberculosis is caused by an obligate airborne pathogen, *Mycobacterium tuberculosis*, particularly by its several human-adapted lineages (1). They represent an intriguing model for the study of evolution due to the lack of horizontal transfer of genetic information, associated with transmission from host to host and absence of an environmental niche (2). The main source of diversity in this pathogen arises from accumulation of mutations, recombination between repetitive DNA, and loss of genetic information through deletions (3).

Specific tuberculosis chemotherapy began with the development of tibione (4) and streptomycin (5) in the middle of the 20th century. The establishment of the complex “first-line therapy” in the 1970s involved the use of four to five effective drugs to prevent the development of resistance (6). In this scheme, rifampicin and isoniazid are particularly potent against *M. tuberculosis*, targeting RNA polymerase and mycolic acid biosynthesis, respectively (7). This treatment regimen is still the cornerstone for susceptible tuberculosis cases (8). However, the global rise of multiple drug resistance and the limited number of antituberculosis drugs (9) require the identification of molecular mechanisms of resistance and the delineation of clinically significant mutations for the personalization of tuberculosis treatment (10,11).

The advent of massive whole genome sequencing since 2013 has provided unprecedented opportunities to study phylogenomics and drug resistance mechanisms (12,13). In addition to genome-wide association studies (GWAS), relying on phenotypic resistance data (14) accounting for homoplasy indexes allows one to identify mutations that emerge independently in different sublineages or clones under selective pressure using only sequence data (12). Additionally, the dN/dS ratio, which quantifies the number of non-synonymous to synonymous mutations in a specific codon or gene, can be used to identify genes under diversifying or purifying selection (14,15). These methods have confirmed many drug resistance mutations and identified new resistance determinants using sets of hundreds to tens of thousands of genomes (14–16). They have also led to the discovery of co-evolved markers that can interact with resistance, increasing minimum inhibitory concentrations (MIC), compensating for the fitness cost of resistance mutations, or improving pathogen tolerance to antibacterial treatments (17,18). Unlike resistance, assessing the virulence and fitness of bacteria is a challenge both *in vitro* and *in vivo* due to the difficulties of phenotype testing (19).

Mutation frequencies of different types are not uniform, even without amino acid or codon selection pressure (20). Substituting C for T in genomes is the most common type of mutation, attributed to spontaneous deamination of cytosine to uracil, which can then result in thymine in offspring (21). The non-uniform frequencies of single nucleotide substitutions shape the spectrum of amino acid substitutions (22) and are included in the Kimura two-parameter model and sequence distance measurements (23). Rarely identified dinucleotide mutations in the same codon (designated further as 2N) are usually considered to evolve sequentially when disctinct species are compared (24,25). The same simplification is used in methods of evolutionary distance calculations, where the frequency of 2N substitutions is set to zero (26).

However, based on genetic code redundancy and inability to obtain particular amino acid substitutions with single nucleotide substitutions the additional value of 2N substitutions as ‘evolutionary shortcuts’ (27) helping to ‘cross the adaption valleys’ (25) could be proposed. Protein structure restrictions favour such complex traits over substitutions obtained by single nucleotide changes in specific cases. Particularly, such mutations could occur in codons encoding the structurally important protein residues, forming the small-molecule binding pockets, or involved into interactions with other proteins, either pathogen, or host nature.

As evidenced by studies on eukaryotic DNA polymerases, 2N substitutions may emerge in a single mutation-repair stage (28,29), reflecting the adaptation of the cell at the amino acid level (30). Analyzing *M. tuberculosis* genomes, we have noticed that these mutations are not rare indeed and could serve as marker of diversifying selection, previously underestimated. We analyzed the frequencies of such mutations from studies on resistance to rifampicin, then, on few recently emerged cases with resistance to novel drugs bedaquiline and linezolid, and finally, we examined 9,941 genomes from the CRyPTIC project for genome-wide analysis of genes under positive selection reflecting both resistance acquisition and host adaptation process.

## 2 Hypothesis: Dinucleotide substitutions in codon reflect a strong selection process at conservative protein sites

### 2.1 Dinucleotide substitutions in *rpoB* gene are frequent in *M. tuberculosis* clinical isolates

Rifampicin has been a cornerstone drug in first-line tuberculosis treatment since the early 1970s, often used in conjunction with isoniazid, ethambutol, and pyrazinamide (6). Resistance mutations to rifampicin occur in the *rpoB* gene encoding the beta subunit of RNA polymerase III (31–33). These mutations are found in four distinct segments known as the rifampicin resistance determining region, which comprises the exit tunnel of the RNA-polymerase complex (34,35) where the drug binding occurs (36).

Many studies of clinically resistant isolates have revealed that the mutation most frequently encountered is S450L (codon 531 in *E. coli* numbering) (37). This mutation is believed to have the least impact on bacterial fitness, and strains carrying it often exhibit extremely high rifampicin MIC. However, many other mutations are consistently found in clinical isolates (38). We decided to analyze the spectra of potential individual amino acid substitutions and reviewed publications starting from a comprehensive TBDreamDB database published in 2009 (39).

Before the era of whole genome sequencing, various methods were used to detect resistance-associated mutations, such as allele-specific or realtime PCR for already known changes in the genome (40) or Sanger sequencing for the identification of new ones (31). We had to exclude data obtained by widely used strip hybridization technology by Hain Lifescience (41) due to partial information loss on specific mutations, detected by the absence of hybridization signal of the amplicon with wild-type probe (42). Some studies using targeted Sanger sequencing lacked raw data and provided only amino acid substitutions, and thus were also excluded from codon substitution pattern analysis. The general data on the proportion of substitutions is shown in Figure 1. We also tried to exclude isolates with multiple mutations that occur in different places in the RpoB to avoid the effects of epistatic interactions, which would require a larger number of isolates to be properly evaluated. We analyzed mutations in both resistant and susceptible isolates because low-level resistance to rifampicin caused by substitutions in codons such as 435 was probably missed. In 2020, the critical rifampicin concentration was lowered from 1 mg/L to 0.5 mg/L for the Bactec MGIT 960 system, and therefore, isolates previously identified as susceptible could indeed have been resistant (43). Additionally, we were interested in analyzing all protein sequence variations that do not significantly affect the activity of RNA polymerase or the pathogen’s adaptation properties.

**Figure 1.**
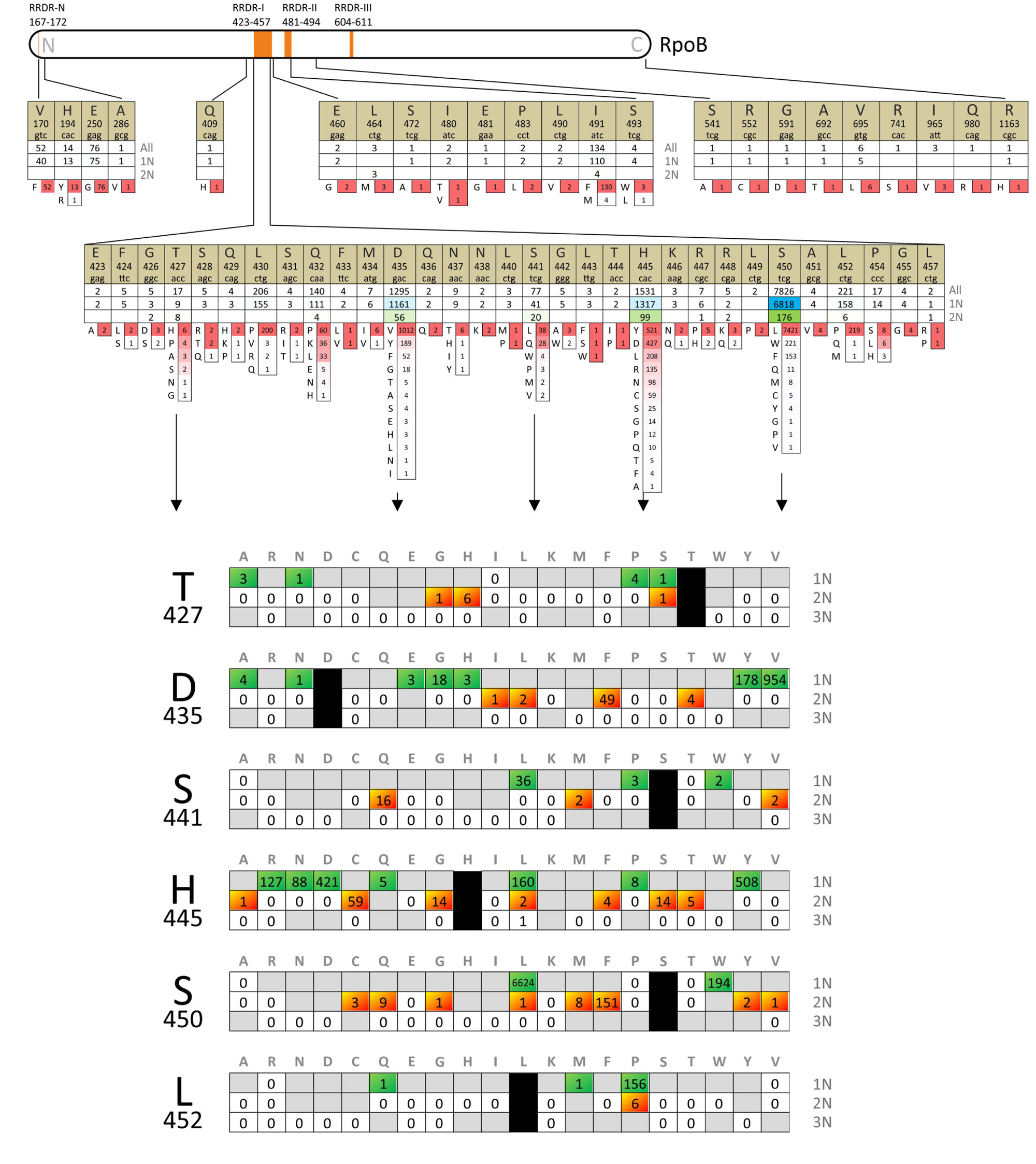
The diversity of *rpoB* mutations obtained from the analysis of 11,730 isolates from the set of 43 studies between 2009 and 2022. For each codon, three numbers are provided: the total number of isolates, the number of isolates with single nucleotide (1N) substitutions, and the number of isolates with dinucleotide substitutions (2N) in one codon. Amino acid substitutions are presented in descending order of frequency, and the corresponding numbers are color-coded from red to white using Excel’s conditional formatting. The frequenctly modified codons are further examined taking into account the number of nucleotide substitutions (1N, 2N, or 3N) required to produce a particular amino acid substitution.

In total, our study included 11,730 isolates from 43 studies (Table S1). More than half of the isolates had mutations in codon 450, followed by codons 445 and 435. Remarkably, a diverse range of amino acid substitution types was observed in all codons. Even at codon 450, where the S450L mutation dominated and was found in 67% of the cases, eight other amino acid substitutions were identified. Seven of these could not be achieved through single nucleotide substitutions, and the total proportion of 2N mutations for this codon was 2.5% (see Figure 1). The most diverse codon was 445, which codes for histidine in the wild-type isolate. Thirteen different amino acids were tolerated by RNA polymerase in this position, and 2N mutations were found in 7% of isolates with an altered 445 codon.

In general, it can be concluded that 2N substitutions provide amino acid diversity that cannot be achieved through 1N mutations (25). However, it is worth noting that, in some cases, the same amino acid substitution could also result from a single nucleotide change, highlighting the inherent stochastic nature of evolution. The frequencies of alternative 2N are two orders of magnitude lower, as is evident from the analysis H445L and L452P mutations (see Figure 1).

In the specific context of drug-induced selective pressure, these substitutions decrease rifampicin binding to RNA polymerase without adversely affecting polymerase activity (44). The wide range of substitution types observed can be attributed to the mechanism of drug action. Rifampicin binds to the DNA/RNA channel at a distance that exceeds 12 angstroms from the active site, effectively obstructing RNA elongation beyond 2-3 nucleotides (34). It could be supposed that the variability would be lower for the drugs that bind directly to more conservative active sites of essential bacterial proteins.

### 2.2 Dinucleotide substitutions, associated with resistance to bedaquiline and linezolid

Bedaquiline and linezolid were introduced into tuberculosis treatment 40 years after rifampicin, in 2014 and 2010, respectively. Since 2014, they have been used in the BPaL combined therapy scheme for drug-resistant TB around the world (45). The safety and efficacy of this scheme led to the redefinition of tuberculosis with extended drug resistance, now defined as resistance to first-line drugs plus fluoroquinolones and bedaquiline or linezolid (46,47). Although acquired drug resistance to these new drugs remains relatively low, there are a growing number of reports of clinical isolates with resistance (48).

The primary mutations selected during bedaquiline treatment emerge in the *rv0678* gene, which encodes the repressor of the efflux operon (49). Loss-of-function mutations in this repressor have a minimal fitness cost and can be distributed throughout the open reading frame (50). Although most of them are frameshift mutations, amino acid substitutions in critical regions are also observed (51). However, the huge number of possibilites to inactivate the gene by frameshifts limits the population frequency of complex and rare substitutions.

In contrast, substitutions in the bedaquiline target, the c-subunit of ATP synthase, which comprises the rotor part, are more conservative and rare, limited by the essentiality of the gene and, probably, by the fitness cost (52). Mutations in the corrresponding *atpE* gene were initially identified *in vitro* and later confirmed in clinical strains of *M. intracellulare* (53) and *M. tuberculosis* (51). To date, there are ten studies reporting 19 strains with substitutions in codons 25, 28, 61, 63, and 66 of *atpE* in clinical strains of patients who received bedaquiline in treatment schemes and developed decreased susceptibility or resistance (51,54–57,52,58–61). The WHO ‘Catalogue of mutations..’ reported eleven isolates with non-synonymous mutations in *atpE*, but these sources of data overlap significantly (38). AtpE I66M and A63P are the most frequent substitutions found in six (56–58,60) and four cases (52,57,60,61), respectively. Additional substitutions in these codons, A63V and I66V were also described (51,54); however, all reported substitutions resulted from a single nucleotide change.

During the study of bedaquiline resistance in tuberculosis cases, we identified an isolate with two simultaneous mutations in *atpE*. This strain had two single nucleotide mutations at different positions, resulting in G25S and D28G amino acid substitutions (51). The simultaneous presence of these two mutations in a single cell was confirmed by whole genome sequencing. We believe such occurrences are less frequent than single-nucleotide mutations, either it was a sequental or simultaneous mutational event.

However, the most intriguing case in the context of the hypothesis was the discovery of bedaquiline resistance caused by a rare dinucleotide mutation in codon 63 (GCA changed to TTA), leading to an A63L substitution (62). No mutations in *rv0678* or other proposed determinants were found in this isolate, yet the minimum inhibitory concentration (MIC) of bedaquiline was more than 2 mg/L (Bactec MGIT 960). A microevolution study of this case revealed a general pattern: the acquisition of the frameshift mutation t141tc in *rv0678* initially occurred in a mixed state, followed by the extinction of the wild-type strain, and ultimately its elimination with simultaneous development of the AtpE substitution. A similar microevolutionary pathway has been proposed based on clinical studies and *in vitro* mutagenesis (63).

Interestingly, some other mycobacterial species are intrinsically resistant to bedaquiline. It was suggested that the presence of methionine in position 63 of AtpE, compared to alanine in *M. tuberculosis*, is responsible for higher MICs in species such as *Mycobacterium flavescens* and *Mycobacterium xenopi* (64,65). Alanine 63 interacts directly with bedaquiline (66) and is substituted with proline or valine in resistant clinical isolates and *in vitro* selected strains (55,67).Both A63P and A63V are the result of 1N substitutions in codon GCA. Other residues, that could be obtained by 1N include E, G, S, and T. The wild-type A, also as P, V and G residues have similar biochemical properties being small, non-polar and hydrophobic. The absence of A63G substitution in clinical strains, resistant to bedaquiline could be explained by the steric properties of glycine, which breaks the alpha helix structure. Dinucleotide substitutions add R, D, Q, I, L, K to the diversity of available amino acids in this position. R, D, Q are hydrophilic, which are unlikely to present of the ATP synthase rotor part exposed to the membrane. Further, K is a positively charged residue, which could have a consequence during the rotation and contact with the stator. Thus, L and I remains the most probable candidates for the 2N substitution of A63 that preserve the function of the enzymatic complex. The substitution of A63 codon for M could be obtained by the change of all three nucleotides in codon (GCA>ATG).

A second example of a rare 2N mutation was observed during a clinical trial in a case of ineffective treatment, resulting in resistance to linezolid. Linezolid inhibits translation by interacting with the ribosome, and resistance mutations are found in distant regions of the 23S rRNA, forming the peptidyl-transferase center. However, most resistant isolates of *M. tuberculosis* carry the substitution C154R in the ribosomal protein RplC, which is typically caused by a single nucleotide change t460c. We identified a resistant isolate with mixed chromatogram data, and Sanger sequencing was unable to distinguish between the simultaneous presence of the nonstandard C154R (CGG codon) and C154S (AGT codon) or two different types of C154R replacements (51). Further whole genome sequencing confirmed the latter, where one subpopulation had the usual t460c mutation, while the other had the TGT codon substituted with AGG by the 2N mutation (Figure 2). The simultaneous presence of two different subpopulations was confirmed by the analysis of subsequent isolates; Even after 15 months, a mixed chromatogram was identified in the clinical isolate obtained from the patient.

**Figure 2.**
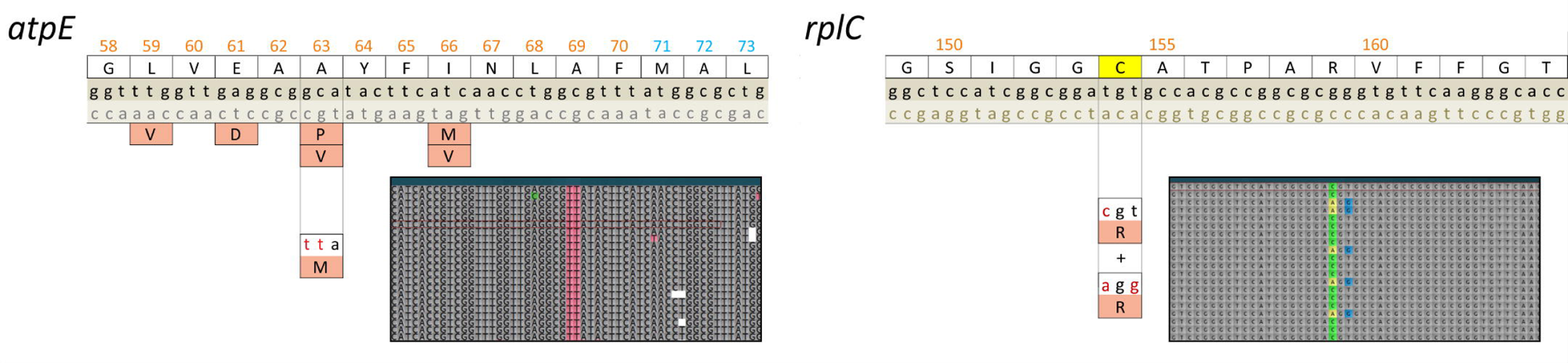
Two cases of dinucleotide substitutions in 63 codon of *atpE* and 154 codon of *rplC* genes identified by the whole genome sequences of the clinical *M. tuberculosis* isolates. Isolates were obtained from the patients receiving the treatment with bedaquiline and linezolid as core drugs, and these substitutions lead to resistance to bedaquiline and linezolid, respectively. The second case contained two subpopulations of *rplC* C154R substituiton – canonical (tgt>cgt) and dinucleotide (tgt>agg) codon substitutions. Screenshots of whole-genome reads (Illumina MiniSeq) aligned to the reference genome of M*. tuberculosi*s H37Rv made in UGENE software.

Both of these cases illustrate the rapid emergence of resistance under the pressure of drug treatment. Similarly to resistance to rifampicin, the 2N mutation in *atpE* expands the diversity of the primary protein structure, while in *rplC*, such a mutation does not lead to novel amino acid substitution compared to the commonly found single nucleotide mutation t460c. Although the number of mutated isolates obtained during and after ineffective treatment is not sufficient to provide strong statistical power for correlation analysis (68), these cases of 2N mutations undeniably provide evidence of ongoing selection in these loci, and directly point to resistance mechanisms. Consequently, it could be proposed that mutations in these codons are clinically relevant and reflect the development of resistance upon the antibacterial treatment.

### 2.3 Dinucleotide substitutions for the prediction of diversifying selection sites in the genome

Although the described examples of bacteria adaptation to drug action unequivocally demonstrate the elevated level of 2N mutations in genetic determinants of resistance, it was interesting to perform a genome-wide analysis of dinucleotide mutation frequencies for the identification of all genes under selection pressure. We assumed that additional resistance determinants and host-pathogen interaction could be found by this approach. For the analysis, we used a data set cosisting of 9,941 of 12,288 isolates that were sequenced and analyzed by the CRYpTIC Consortium (69,70). The remaining 2,347 had an ambiguous description of amino acid substitutions and were omitted.

Two main approaches were used for the analysis of mutation frequencies: allele counting and homoplasy counting (71). Although the former uses frequencies observed in the population and does not require intensive calculations of phylogeny and ancestor reconstruction of mutation events, it has significant disadvantages in its application to *M. tuberculosis*. The clonal nature of the pathogen led to complete linkage disequilibrium and population stratification, which could result in false associations obtained by uncorrected allele counting methods (69).

The homoplasy count derived from the number of independent events in different branches (12) could be a signal for positive selection (72). Following this method, we constructed the phylogenetic tree using mutations from all genes, excluding highly repetitive PE, PPE, PE-PGRS genes and insertion elements (Figure S1). The reliability of the phylogenetic tree was verified by analyzing the distribution and frequency of selected lineage-specific SNPs (73).

The maximal parsimony approach was applied to the set of all variations at a specific codon. All mutations were mapped on the tree and traced by joining neighbors with this mutation and mapping it to the common ancestor. It was supposed that back mutations could have occurred if it allowed the joining of branches with mutations split by smaller branches with the wild-type codon. Thus, the final number of all events was minimized for each codon across the whole tree.

As an example, the mutation E112K in the *rv0078A* gene (codon change GAG>AAG, nucleotide substitution c87468t), which could be used as a phylogenetic marker of lineage 2 (73), was present in 3,838 of 9,941 isolates. Indeed, they all belong to the lineage 2 branch (*n* = 3,842 isolates) of the constructed *M. tuberculosis* phylogenetic tree; however, four isolates in this branch did not have this mutation. The phylogeny-adjusted frequency of independent events in this case for the entire population was equal to 23. The introduction of the maximal parsimony approach, based on the proposal that in these four isolates the reverse substitution occurred recently, allowed the decrease in homoplasy counts to 2 for direct and reverse mutations (four genomes with ‘wild-type’ Lys at 112 constitute one branch). Therefore, the direct mutation was traced to the common ancestor of lineage 2.

In such an approach, the consequent 1N mutations leading to final 2N substitution were treated as separate, thus underestimating the final diversity of codon substitutions and decreasing the numbers of 2N events. However, counting the second mutation as 2N will lead to an overestimation due to cases where the first mutation was specific to the branch, for example, the whole lineage 2. Thus, the analysis was limited to the calculation of simultaneous 2N substitutions, providing a more specific but less sensitive result in terms of codon diversity at particular position.

The resulting number of events for 275,531 codons in 3,680 genes analyzed were the following: 142,321 events of synonymous substitutions, 252,702 events of 1N substitutions and 1,645 events of 2N substitutions. The latter occurred in 862 codons in the genome, with a maximum value of 350 for the substitution of *cyp138* P114F. Excluding the genes with single type of 2N substitutions results in 251 codons in 97 genes. The comparison of the number of codons with 2N substitutions and 2N events for each gene is shown in Figure 3. The two most important genes, that cause resistance, *rpoB* and *katG*, have elevated parameters and are clearly distinguishable from the entire set of genes.

**Figure 3.**
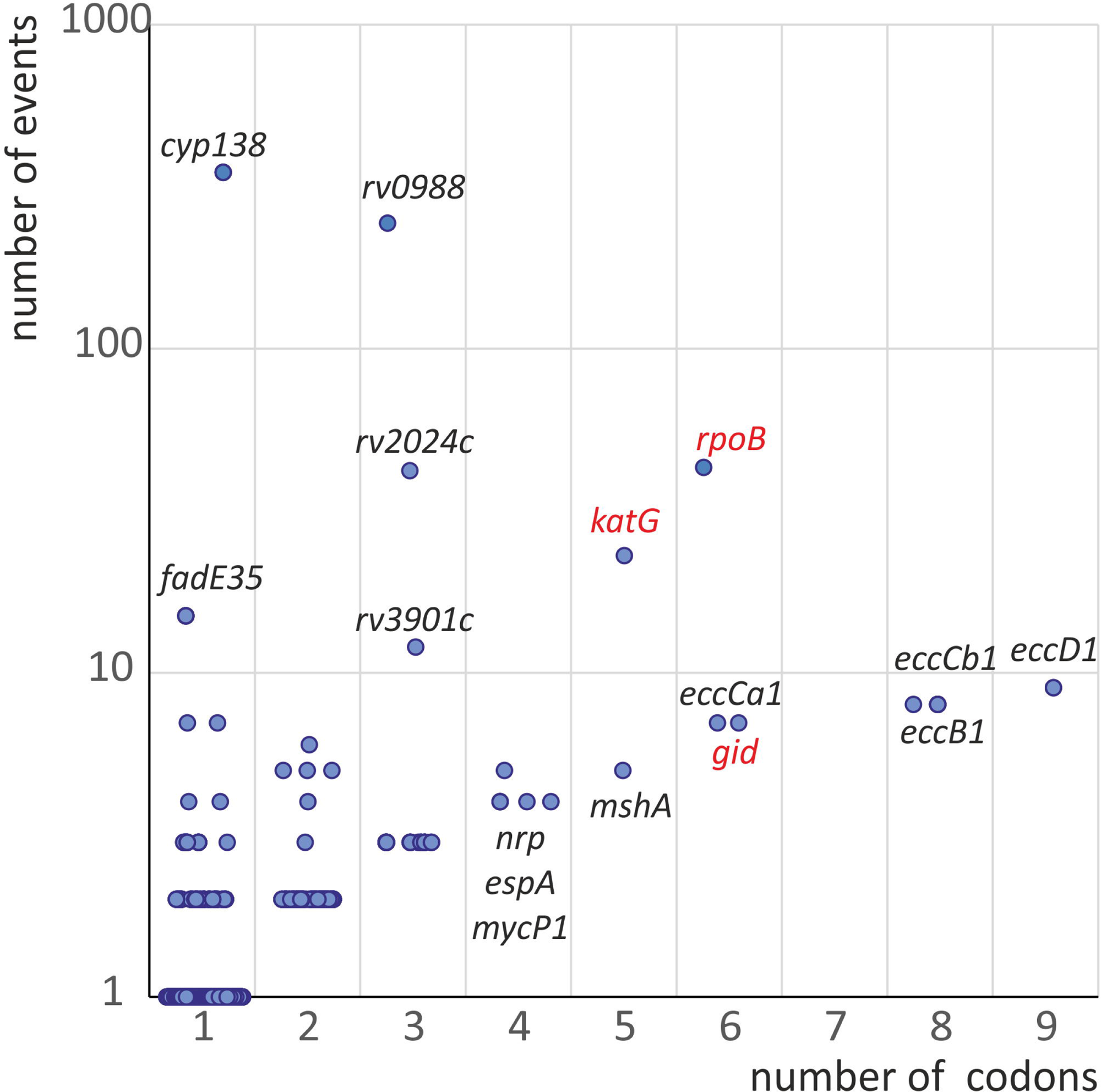
The variability of phylogenetically-adjusted frequencies of 2N substitutions and number of affected codons in *M. tuberculosis* population (CRYpTIC study). Known drug resistance determinants are marked with red.

Splitting of the counts of nonsynonymous mutations into 1N and 2N mutations allowed the introduction of two metrics, the ratio of dinucleotide to single-nucleotide nonsynonymous substitutions d2N/d1N, and ratio of dinucleotide to synonymous substitutions d2N/dS. In the commonly used GWAS approach, the comparison of dN/dS in resistant and susceptible sets of isolates is performed for identification of resistance determinants or genetic traits that co-evolve with resistance (69). We used another approach for identification of all genes under selection by the use of codon-based models of evolution and estimation of the difference between measured and expected frequencies of codon substitutions (74,75).

The codon substitution matrix for the Monte-Carlo modeling was derived from the whole dataset of phylogeny-adjusted events. Additional correction was introduced by proportional use of all variants of the particular codon in the whole set of isolates as the starting codon in modeling. Otherwise, lineage-specific codons with high prevalence in the genome set would not add to the calculated substitution frequencies. The difference between observed and estimated frequencies allows the identification of any genomic traits that are putatively under selection (76). The statistical approach for the identification of significant traits was analogous to that of the widely used ratio of non-synonymous to synonymous substitutions dN/dS (69). Therefore, three values for genes were analyzed and compared simultaneously presented as Manhattan plots (Figure 4), two for 2N substitution statistics and conventional dN/dS was used as reference. The significance threshold of - log(*P* value) was estimated by Bonferroni correction (77).

**Figure 4.**
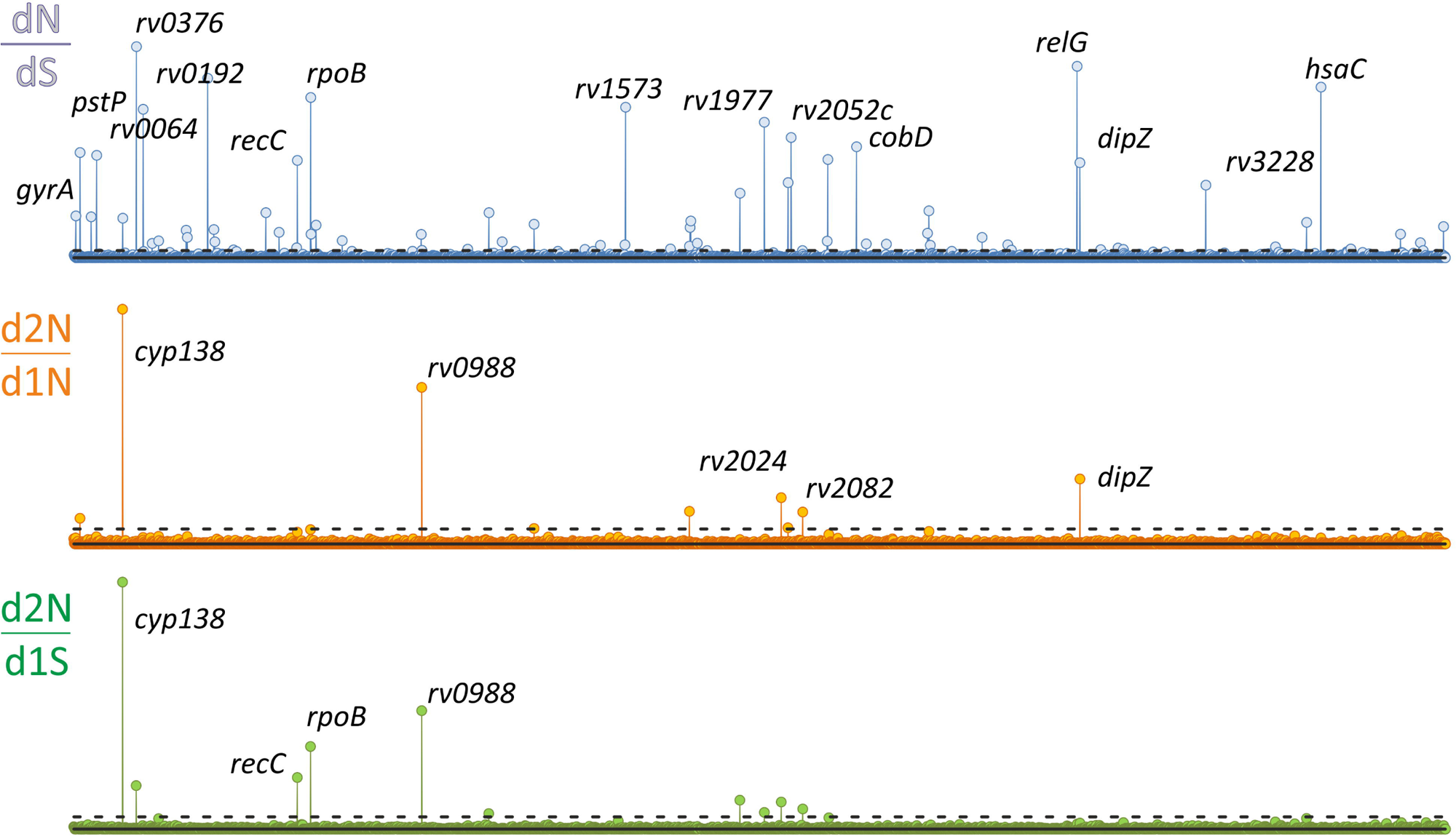
Genome-wide study of traits under selection from the analysis of 9,941 *M. tuberculosis* genomes of the CRYpTIC study. Manhattan plots for the three ratios were used: total non-synonymous to synonymous (dN/dS), dinucleotide to synonymous (d2N/dS), and dinucleotide to single nucleotide substitutions (d2N/d1N) are plotted along the chromosome. The statistical significance identified by the Bonferroni correction are shown with dotted lines.

The phylogeny adjusted counting of single-nucleotide mutations of different type and the comparison between genes resulted in distinct statistical signals for 68 genes, including the main resistant detrminants *rpoB*, *katG*, *rpsL*, *pncA*, *gid, embB*, and *ethA* (Table S2). Fourteen genes had a statistically significant change, either increase or decrease, in the dinucleotide events statistics, including the resistance determinants *rpoB*, *katG*, and *pncA*. Two genes, *rv2024c* and *rv2082*, with low *p*-values of d2N/d1N and d2N/dS, were not significant by dN/dS analysis (Table 1).

**Table 1.**
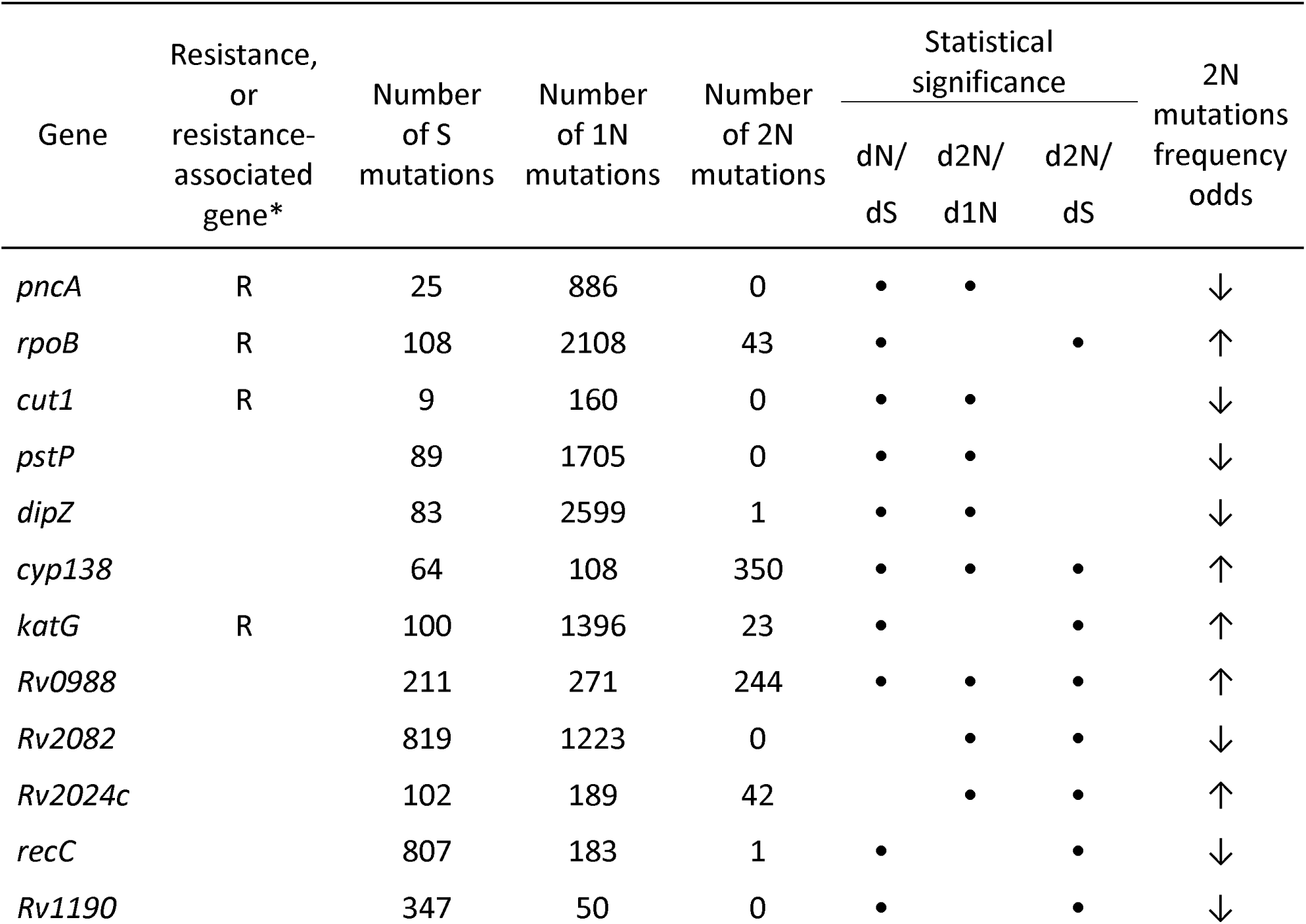
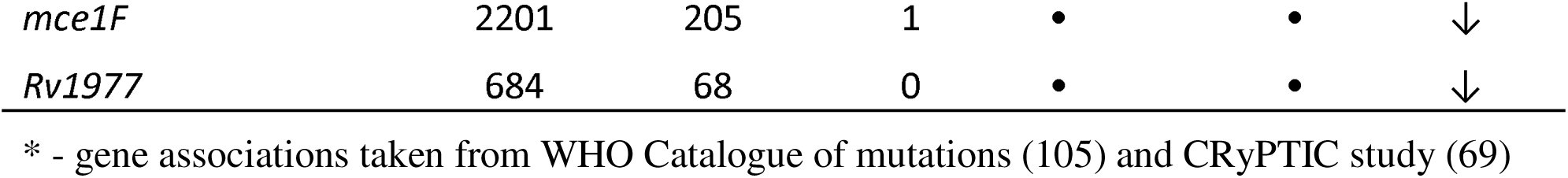
Top significant genes identified by the elevated 2N events in single codon.

The statistically significant increase in 2N substitutions was observed for five genes only (Table 1). The *rpoB* gene had the most diverse mutation types and the largest number of isolates due to its role in resistance to rifampicin. Eighty-seven isolates with twelve 2N substitutions in six codons were found. Homoplasy counts of 2N mutations in the whole gene were 43, while that of 1N mutations was 2,216 (3%). Several types of dinucleotide substitutions were identified in codons 435, 445, and 450, similar to the pattern described in the Section 2.1. An additional new 2N mutation, I542K, was also found.

In addition to *rpoB*, the main isoniazid resistance *katG* gene was found to be significant by dN/dS and d2N/dS ratios. The most prevalent S315T mutation found in resistant isolates is obtained by single nucleotide G>C substitution (codon AGC is changed to ACC). In this set of isolates, three other dinucleotide substitutions were identified thaqt led to the same amino acid change: AGC codon was changed to ACA, ACG, or ACT.

The favored use of particular amino acids in drug-binding pockets with low fitness-cost could explain this observation for resistance genes *rpoB* and *katG*. Other genes associated with resistance to first- and second-line drugs in *M. tuberculosis*, such as *rpsL*, *embB*, *gyrA, gid*, and *ethA*, were significant only by dN/dS analysis.

Two more genes, that were not previosly linked with resistance, had significant statistical signals for 2N substitutions, *cyp138* and *rv0988*. Both amino acid substitutions, *cyp138* P114F (CCT > TTT) and *rv0988* L191A (CTG > GCG) cannot be obtained by single nucleotide substitution, confirming the importance of particular amino acid residues at these positions.

The predicted cytochrome P450 *cyp138* is involved in pH tolerance mediated by MTS1338 sRNA regulation (78). Cytochromes P450s Cyp138 join metabolism, virulence, and susceptibility to antibiotics by altering cell wall composition (79). The gene *rv0988* is a subunit of the ABC transporter (80) and negatively regulated in the presence of toxic levels of CuCl2 (81).

Both *cyp138*(P114F) and *rv0988*(L191A) emerged in many unrelated strains of the *M. tuberculosis* lineage 2 (East-Asian) and were absent in all other lineages: lineage l (Indo-Oceanic), lineage 3 (East-African-Indian), lineage 4 (Euro-American). Cyp138 (P114F) is present in 3,154 of 3,527 isolates of lineage 2.1. Such lineage-specific distribution differ from the distributions of resistance-associated variants, which are present in all human-adapted lineages – L1, L2, L3, L4, which is cause by the common therapy used in different regions of the world. Thus, it could be supposed that mutations in *cyp138* and *rv0988* are related to the pathogen adaptation to hist pressures from immune system and antituberculosis drug use.Alternatively, these traits could be responsible for the evolutionary success of Lineage 2, which was widely discussed in the phylogeography studies on tuberculosis (82).

Using the maximal parsimony approach, we suppose that P114F occurred only once in the common ancestor of 2.1, and then, in 373 isolates the reverse substitution F114P emerged (Figure 5). The phylogeny adjusted frequency of the reverse mutation is 349, resulting in 350 total events in the codon for this set of isolates. Alternatively, single-stage accumulation of P114F in the 3154 genomes demands 1130 mutational events.

**Figure 5.**
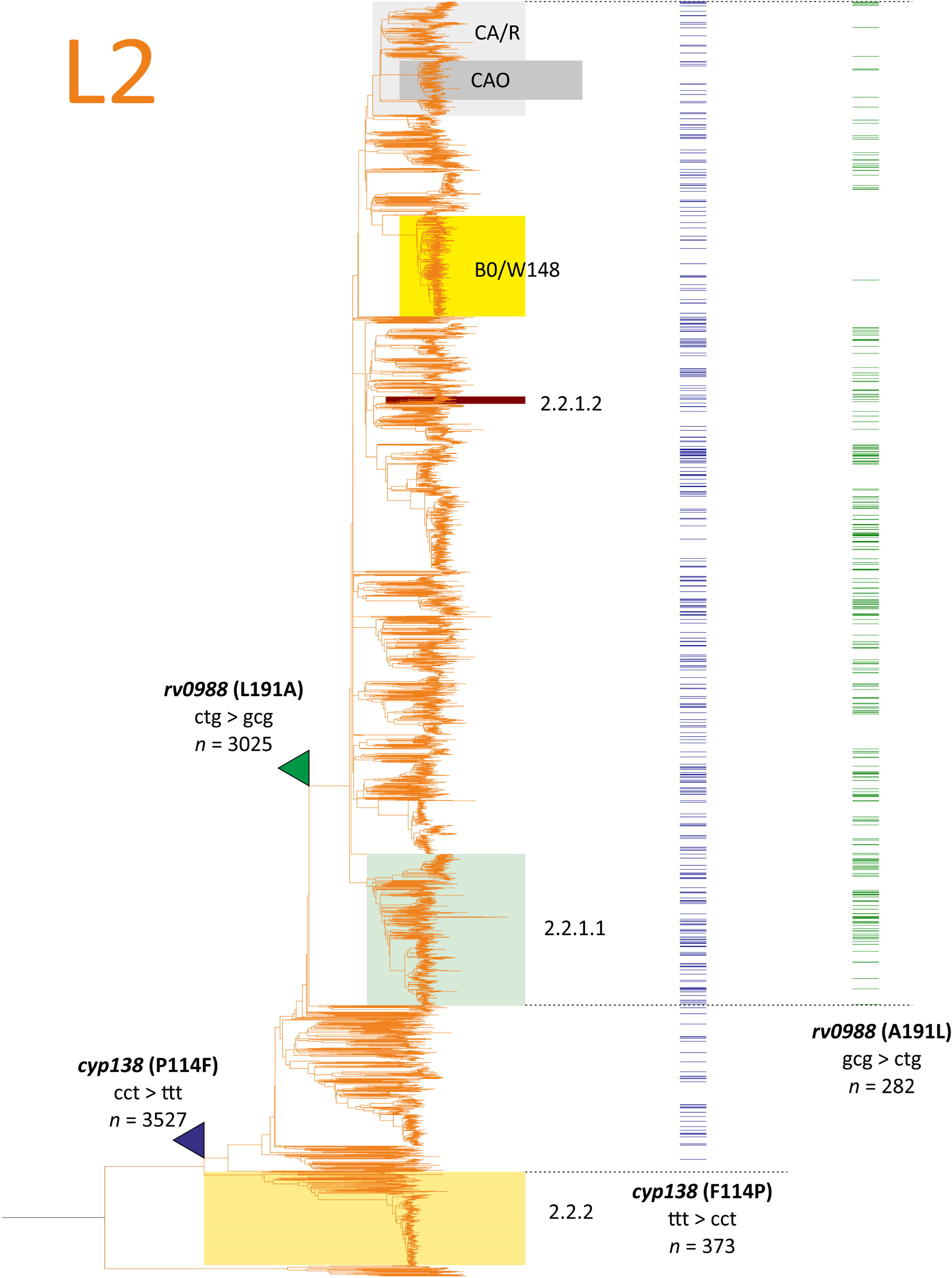
The maximal parsimony model of selection of lineage 2 specific 2N substitutions *cyp138*(P114F) and *rv0988*(L191A) from the analysis of 9,941 *M. tuberculosis* genomes of the CRYpTIC study. The branches where probable direct substitutions occurred are pointed with color triangles, reversions are shown as distributions in terminal branches (isolate genomes). Number of isolates with mutations in the whole CRYpTIC set are given unadjusted.

The emergence of *cyp138*(P114F) has recently been described as having evolved under the pressure of HIV-1 coinfection (83). However, taking into account the model of sequential forward and reverse mutations in Lineage 2 strains, the opposite situation could be supposed: the revertion F114P is occurres under the pressure of immunocompetent HIV-naïve host.

The same considerations are also relevant for *rv0988*(L191A): forward substitution in the common ancestor of a smaller sublineage within 2.1 and reversion to Leu in many unrelated strains, which is common for all other lineages (Figure 5). The phylogeny-adjusted frequency in the absence of reversion hypothesis was 742, compared to 241 events of reverse substitutions. Interestingly, the distribution of A191L revertion is not unified in L2 sublineages. For example, the Beijing BO/W148 genotype, which is a successive clone spreading in Russia (84), has a low frequency of reversions, while some other branches are nearly completely reverted (Figure 5). Such a non-random purge of probably deleterious polymorpisms (85) demands further studies on correlation with geographical localization of clnical isolates and impact on virulence and pathogenicity.

Contrary to the previous, the substitutons N508T and C514L in *rv2024c* could be achieved also by simple 1N substitutions in corresponding codons, however, the codon usage in these cases is significantly lower compared to obtained by observed 2N substitutions. Thus, the observed AAT>ACG for N508T changes frequency from 5.3 to 35.1, while the codon Thr codon ACT has frequency of 3.7 (86). Similarly, for the C154L codon usage frequency increased from 8.1 to 50.4, while the alternative Leu CTA codon, which could be obtained by 1N substitution, has frequency of 4.8 (86). The exceptional use of 2N substitutions in unrelated strains also points to the preference for particular amino acid at these positions of the protein. Rv2024 has an unknown function but showed diversity in parallel and sequential isolates during the within-host microevolution analysis (87) and is believed to have a virulence-associated function, since it is was found to be induced in mouse lungs (88).

Two more classes of genes could be identified among those with a low number of 2N substitutions: under positive selection (dN>>dS) and under purifying selection (dS>>dN). A high number of non-synonymous substitutions were identified for *pncA*, *cut*, *pstP*, *dipZ*, and *Rv2082* in the top-scored list of genes (Table 1). Mutations in *pncA* that lead to the loss of function of the gene are selected during pyrazinamide treatment and lead to resistance. Such mutations are frequent, distributed along the open reading frame, and include frameshifts. Therefore, the low number of 2N substitutions is not an unexpected phenomenon, also as for the *rv0678c* gene alterations leading to the resistance to bedaquiline. In addition to selection for loss of function, genes with high 1N and low 2N counts could encode proteins that are tolerant to substitutions, or the possible changes with low fitness cost are achieved by 1N codon substitutions.

Genes with a high number of synonymous substitutions have low counts of 1N and 2N substitutions, and the statistical significance of the d2N/dS ratio is the direct consequence of the significance of dN/dS ratio.

## 3 Discussion

Massive whole genome sequencing has allowed unprecedented possibilities for studying evolution and cellular mechanisms, including drug resistance. Various approaches have been developed, such as GWAS (69), dN/dS (13,85,15) and PhyC (12), which are mostly based on estimating mutation frequencies at specific genomic locations and correlating them with phenotypic characteristics. These methods involve calculating the ratio of non-synonymous substitutions to synonymous substitutions and assessing homoplasy, where the same type of mutation is present in different sublineages of *Mycobacterium tuberculosis*. Since there is no known horizontal gene transfer and a known environmental niche, mutations in different lineages and sublineages occur independently, reflecting selective pressures.

Bacterial evolution pathways are highly diverse and, in the case of tuberculosis, transmission and survival within a host have shaped the pathogen’s adaptation over thousands of years (3). The recent introduction of antibacterial drug treatment, starting with drugs such as streptomycin and tibione, has become a dominant force driving evolution of the pathogen. In this context, diversifying selection has led to changes in amino acids that interact with drugs in their target proteins (89). However, it should be noted that in cases of ineffective treatment and the transmission of resistant isolates within a population, we may observe purifying selection, where mutations that result in fitness costs are eliminated from the population (90). On the other hand, fitness-compensating mutations may emerge, representing another form of diversifying selection driven by forces distinct from direct drug action (91).

The development of drug resistance is a time-limited event, whereas host pressures drive the ongoing competitive evolution of both the pathogen and the host (89). This phenomenon, often referred to as the “Red Queen Effect” (92,93), influences various virulence factors in *Mycobacterium tuberculosis*. These factors enable the pathogen to survive within macrophages by suppressing apoptosis, disseminate throughout the host, utilize diverse nutrient sources, and manipulate host cell processes (94).

Both directions of evolution are highly interconnected, since resistance selection drives the adaptation process through mutagenesis. Recent findings indicate that selection for resistance to rifampicin, for example, leads to an increase in the genome-wide mutation rate, thus accelerating evolution (95).

The diversity of mutations in a population results from the combined effects of chemical mutagenesis, repair mechanisms, and possible selection preventing the essential functions of the cell, ensuring the adaptability and low fitness cost. Other, less measurable or estimated, pressure forces originate from the secondary structure of RNA, its stability, cross-interactions between different RNA molecules, transcription and translation efficiencies, and more (96,97).

From a protein perspective, the structural and functionally tolerated variability of amino acids at a specific position cannot be achieved by single nucleotide changes alone. While a single nucleotide change at a particular codon can result in approximately seven amino acid substitutions, more complex dinucleotide and trinucleotide mutations can generate all the range of peptide sequences, albeit with lower probability. As a result, the frequencies of 2N mutations leading to beneficial amino acid substitutions that cannot be obtained by 1N substitution in codon tend to be higher compared to the baseline frequency. Another driving forces for preferral use of 2N substitutions could come from codon usage frequency, secondary mRNA structure, and antisense mRNA interactions. In this case, the same amino acid substitution could be achieved also by single nucleotide substitution.

We observed both types of 2N substitutions in *M. tuberculosis* genomes; however, the total number of mutational events was low. The described approach could provide much informative results if the significantly larger set of genomes will be analyzed. Nowadays, more that 200,000 genomes of *M. tuberculosis* is available in NCBI SRA database (*n*=219,660, databased accessed on November, 23, 2024). Since no phenotype data is needed for the described approach, more genetic traits significant for the pathogen adaptation would be identified upon the expanding the analyzed set of genomes. We observed many genes with altered frequency of 2N substitutions, that did not achieved the desired statistical significance threshold. They include already known genes that are associated with resistance, or co-evolved with the emergence of resistance: *gyrA*, *ethA*, *rpoC*. The statistical significance of 2N substitutions in these genes will be improved with the growing number of isolates, as well as for other important unidentified genes and codons. However, the inclusion of mutational data for hundreds of thousands of genomes compared to ten thousand used in this study will require much more intensive data processing and advanced computing facilities.

While the current knowledge about tuberculosis drug-resistance mechanisms was summarized in two issues of WHO’s ‘Catalogue of mutations’, virulence-associated traits remains to be obscure in general. An important result of the tuberculosis evolution studies was the description of ‘successful’ clones or sublineages rapidly spreading in population (84). Such clones are associated with resistance, virulence and poor outcomes. The developed molecular methods for the identification of such important genotypes, most probably, have its main clinical significance in preventing of superinfection in hospitals. However, we do not know, which particular mutation, or complex of mutations were responsible for the evolutionary success of these sublineages, while the difference between pathogenic properties of different genotypes has been observed in model experiments (98).

Experiments with live *M. tuberculosis* are not easily performed due to the safety considerations, low growth rate, and limited number of genetic tools. Even not every resistance-associated variant was verified by its introduction into wild-type strain and phenotype testing of the recombinant clone. The model organism *M. smegmatis* is not suitable for virulence estimation, and the mouse model of tuberculosis infection does not reflects all the aspects of human infection (99). Thus, statistical apporaches of massive genome analysis remains the main source of information of resistance, fitness-compensation and virulence traits of the *M. tuberculosis* genome.

In a massive genome-wide analysis using the dinucleotide statistics we observed the strong signal of selection in two genes not associated with resistance, *cyp138* and *rv0988*. The frequencies of substitutions were elevated in two sublineages of lineage 2. Since we used the hypothesis of possible reversions and maximal parsimony approach minimizing the number of events, we supposed that the direct substitution in the common ancestor occurred first, and then many back substitutions took place reverting to the aminoacid that is common to all other lineages in this position. The reversion hypothesis have some limitations. First, only single reversion was allowed, which we suppose to be true due to low mutation rate in *M. tuberculosis*. Second, while the reversion is likely for the case of E112K substitution in the *rv0078A* gene, which were present in 3,838 isolates of 3,842 lineage 2 isolates, the cases of substitutions in *cyp138* and *rv0988* genes have much more comparable counts with or without reversions. However, the number of mutational events without the reversions would have been three times higher, and if we used such higher counts of events, the confimation of 2N substitutions importance in these cases would have been more statistically significant. By minimizing the events number we thus used more strict approach that could have missed some traits, but the provided result is more specific.

The result obtained in this study provide some clues to some novel mechanisms behind the adaptation of the *M. tuberculosis*. We observed not only the preferral use of complex dinucleotide mutations at particular codons, but also the lineage-specific selection of mutations, which is most probably caused by some host pressure, either from the host immune response or drug use. The frequency of Rv0988 A191L amino acid substitution is extremely low in ‘succesfull’ sublineages Beijing B0/W148 (84) and Central Asia/Russian (100) compared to other Beijing clones. Future introduction of such traits into diagnostics could allow the personalized therapy accounting the pathogenic properties, and the rapid identification of new epidemiologically significant lineages at their earliest stages of spreading in the population. Moreover, the conserved nature of substitutions caused by dinucleotide mutations give hope for the development of virulence-targeted therapy aimed at these particular residues (101).

It should be noted however, that there are few other alternative possible mechanisms explaining the observed mutational events elevation instead of natural selection objected by the interaction with host immune system (19). While we could reject the 2N substitutions neutrality with high confidence due to the statistics, the nature of substitutions, and tha absence of horizontal transfer, such nonrandom distributions could be caused by the coevolution with human population. Besides the adaptation to the particular subpopulations, the human densities and immigraton could shape the genetics of the pathogen (94). The lineage specific homoplasy events could also be the result of HIV coinfection or variability of vaccination strategies used in different parts of the world (90,94). Dinucleotide and other more complex mutation occurred in all branches of life, providing the additional diversity for survival and adaptation. To confirm the universality of the finding, and the possibility to apply the approach not only to clonal *M. tuberculosis*, we performed the same analysis for 1,164 genomes of

*N. gonorrhoeae* (supplementary data). In this set of genomes, 2N substitutions constituted 1.1% of all nucleotide substitutions. Similarly to the *M. tuberculosis* genomes, the fraction of genes with altered 2N substituitions took only a small part of the whole set of genes with high number of all point mutations, calculated from dN/dS statistics. Most frequently, mutations in genes, encoding monomers of the *Neisseria* type IV pili were identified. Type IV pili of *Neisseria* encoded by a higly variable genes set which diversity is driven by intergenome recombination, and is crucial for virulence. Pili are involved in surface adhesion, bacterial cell aggregation (102), they are recognized by specific antibodies. and alter host signalling (103). Thus, particular changes of specific residues are not unexpected, and are limited by its functional involvement (102).

In conclusion, the distinct distribution of dinucleotide mutations reveals unique insights into amino acid selection preferences. Genes exhibiting frequent 2N mutations are often pivotal for critical interactions with other molecules or proteins either pathogen, or host nature, playing a fundamental role in the pathogen’s survival. The elevated occurrence of 2N mutations signifies diversifying selection, encompassing drug resistance mechanisms and factors associated with host invasion and survival. Exploring these interactions, virulence determinants or their counterparts, presents a promising avenue for innovative therapeutic strategies (104). However, it is crucial to acknowledge that the substantial diversity and rapid evolution within these genes can pose challenges to the long-term effectiveness of such therapies. Therefore, an approach that focuses on more conservative regions of the identified targets might be preferable in drug development.

## 5 Author Contributions

DZ – conceptualization, data curation and formal analysis (section 2.3), visualization, writing; AU - data curation and formal analysis (sections 2.1 and 2.2), writing

## 7 Data Availability Statement

The datasets analyzed and generated for this study and the developed scripts can be found in the Zenodo.org at https://doi.org/10.5281/zenodo.15367741.

