## Supplementary methods for "Dinucleotide codon mutations as a marker of diversifying selection in *M. tuberculosis*"

#### Isolates with bedaquiline and linezolid resistance

The isolates analysed in the study were from our previous reports (Zimenkov, 2017, Peretokina, 2019, Ushtanit, 2020). Isolates were obtained from patients who received bedaquiline and linezolid as the core drugs during treatment. All isolates were investigated by phenotypic bedaquiline susceptibility testing on Middlebrook 7H11 agar and Bactec MGIT 960 liquid media. Resistance to linezolid (Sigma-Aldrich Co., St. Louis, MO, USA) resistance was determined using a MGIT 960 (critical concentration of 1mg/L).

| Patient Iq | *rv0678* – t274ta*  AtpE – G25S* D28G* | MIC Bdq:  0.25 mg/L (agar)  1 mg/L (liquid) |
| --- | --- | --- |
| Patient Jk | AtpE – A63L (gca🡪tta) | MIC Bdq:  >1 mg/L (agar)  4 mg/L (liquid) |
| Patient Bg | RplC – C154R (tgt🡪cgt*/agg*) | Lzd - R |

* - heteroresistant state.

Whole genome sequencing was additionally performed for isolates from patients Iq and Bg. The isolate from patient Jk was sequenced previously (Ushtanit, 2020, accession SRR16168839).

Genomic DNA was extracted using the Gentra Puregene Yeast/Bact. Kit (QIAGEN cat no. 158567). Briefly, 0.25 mL of culture suspension aliquots were centrifuged at 13500 rpm for 5 min. The pellet was washed and treated with Puregene cell lysis buffer and Proteinase K. The DNA was washed with ethanol and resuspended using a a hydration solution. The DNA concentration was measured with a Qubit HS DNA fluorometer (Life Technologies, Carlsbad, CA, USA). Whole genome sequencing was performed using the MiniSeq platform (Illumina, San Diego, CA, USA).

Sequencing data in FastQ format was analysed using the internal pipeline. The reads were trimmed using the Trimmomatic tool (Bolger, 2014), mapped to the reference genome of *M. tuberculosis* (GenBank accession NC_000962.3) (Cole, 1998) with BWA-MEM (Li, 2010) and refined using BamLeftAlign [59]. Variant calling was performed using FreeBayes (Garrison, 2012) and filtered with the VCFlib toolkit. Variant annotation was performed using SnpEff (Cingolani, 2012).

The raw sequencing data was submitted to the Sequence Read Archive of the National Centre for Biotechnology Information (accession number PRJNA768108).

#### CRyPTIC database retrieval and analysis

Molecular and phenotypic data were obtained from the official CRyPRTIC FTP server [http://ftp.ebi.ac.uk/pub/ databases/cryptic/](http://ftp.ebi.ac.uk/pub/%20databases/cryptic/). The following files were downloaded from the subdirectory /release_june2022/reproducibility/data_tables/cryptic-analysis-group (accessed on July, 2023):

- VARIANTS.csv, containing data on nucleotide mutations,
- MUTATIONS.csv, containing data on amino acid substitutions and codon changes, and
- CRyPTIC_reuse_table_20211019.csv, which contained phenotypic data for 12288 isolates.

Data processing and analysis was performed using custom Python scripts. First, all amino acid substitutions and SNPs for 12288 isolates were extracted from the VARIANTS and MUTATIONS tables into separate tables.

Highly repetitive PE/PPE genes, insertion elements, and phages were excluded from the analysis. The 3680 genes of total length 3.62 Mbases comprised 82% of *M. tuberculosis* H37Rv genome (NC_000962.3).

Two thousand three hundred and eighty-eight isolates were excluded from the study due to the presence of mixed and unknown mutations marked as ‘x’ or ‘o’ in the annotation. Mutations of the final set of 9941 isolates were remapped onto *M. canetti* genome (NC_015848.1). The list of amino acid substitutions was used to build the Python *numpy* and *gmpy2* tables with binary data on the presence of particular substitutions for all isolates.

The phylogenetic tree was built using the Nearest-Neighbout approach (Saitou, 1987) and the MEGA 11 software (Tamura, 2021). The robustness of tree was checked using lineage-specific SNPs (Coll, 2014). Isolates belonged to four main lineages – Indo-Oceanic (L1), East-African-Indian (L3), East-Asian (L2), and Euro-American (L4) (Figure S1).

Each genomic mutation was mapped to the phylogeny using parsimony approach. First, the lists of nearest neighbours and descendants for each terminal branch corresponding to isolate were obtained. If some mutation was absent in the analyzing isolate, but were present in the neighbours and ancestors, we supposed that the both the common and reverse mutation occurred in this case, and the list of mutations was updated to include these two steps in the analyzed isolate. The same approach was used for the same gene and codon mutation of different type – such mutations were splitted into common ancestral mutation plus change from ancestral to isolate-specific mutation (Figure S1).

These corrections of mutation lists allowed to map mutations along the phylogenetic tree and identify the nodes where mutation emerged. The initial frequencies of mutations (number of isolates with mutations) were corrected to number of nodes with mutations, which is drastaclly lower for the most of mutations (Table S2). In total, 325,419 different mutations were found in this set of isolates. The initial and phylogeny-adjusted numbers of mutation events in population were equal to 42,379,768 and 449,223, respectively. Adjusted frequencies of 3-nucleotide substitutions in one codon and dinucleotide substitutions, leading to synonymous amino acid substitutions (Arg, Leu, Ser), were as alow as 42 and 97, and were omitted from the analysis.

For the statistical analysis of mutational events the Fisher exact test was used as in widely used GWAS approach, also designated as dN/dS and branch-site test (Farhat, 2013; Chiner-Oms, 2022; Liu, 2022). Breakpoint values of *p*-values (-log_10_P-value) were selected on the basis of Bonferroni correction, three deviations from the the expected value using the analysis of distributions, or QQ-plotting (Taboga, 2021). The most strict values were used.

Three main values were analyzed – the conventional ratio of nonsynonymous to synonymous substitutions dN/dS, ratio of dinucleotide to single-nucleotude nonsynonymous substitutions d2N/d1N, and ratio of dinucleotide to synonymous substitutions d2N/dS. The results of the latter two approaches had a greater priority for the further analysis, particularly in the case if the mutation event was insignificant by dN/dS approach.

Both source frequencies (number of isolates with and without the mutation) and phylogeny-adjusted frequencies were used in several statistical calculations.

First, source frequencies were splitted based on resistance profile of the isolates and associations of mutations of different types with resistance were analyzed by conventional Fisher exact test. From the whole set of isolates 3695 were resistant to at least rifampicin, and 5429 to any of the drugs analysed by CRYpTIC consortium (rifampicin, rifabutin, isoniazid, moxifloxacin, levofloxacin, ethambutol, ethionamide, kanamycin, amikacin, clofazimine, bedaquiline, linezolid, delamanid). Manhattan plots of dN/dS, d2N/d1N, and d2N/dS derived values are shown on Figure S2.

Another approach was based on pairwise comparison of mutation frequencies in different genes with normalization per gene length. To construct the matrix of -log_10_(P-values) for each pair of genes dN and dS values for gene *i* were compared to dN and dS values of the gene *j*. Average values for each gene were calculated at each row of the matrix and plotted (Figure S3).Two calculations were performed using initial and phylogeny-adjusted numbers of mutations (allele and homoplasy counting).

The fourth approach was the estimation of the difference between phylogenetically adjusted mutational events and expected values at the level of individual codon. To obtain the null hypothesis for comparison the codon substitution matrix (20 x 20) was obtained from the total number of phylogeny-adjusted codon substitution events (n=449,223). Then, for each codon expected values of synonymous, non-synonymous caused by single- and dinucleotide substitutions were estimated from the matrix using Monte-Carlo method. Number of events were balanced with the total number of observed mutations in particular codon. For simplification of data presentation, maximal *p*-values for each gene are provided on Manhattan plot (Figure S4) and in a table format (Table S3).

#### Correction of annotation

In the article by Koch et al., 2017, cited in the Section 2.3, the annotation of mutation in *cyp138* gene was not correct due to the simultaneous presence of two SNVs in one codon. They annotated c163705t as P114S, while the correct substitution is cc163705tt, leading to amino acid substitution P114F. Genomes from this study were partially reanalyzed, and at least the following genomes with accessions ERR1633777, ERR1633797 have both SNVs.

ERR1633775, ERR1633776, ERR1633778, ERR1633779, ERR1633780, ERR1633781, ERR1633782, ERR1633783, ERR1633784, ERR1633785, ERR1633786, ERR1633787, ERR1633788, ERR1633789, ERR1633790, ERR1633791, ERR1633792, ERR1633793, ERR1633794, ERR1633795, ERR1633796 were wild-type.

#### Neisseria genomes analysis

The set of *Neisseria gonorrhoeae* genomes were analyzed using the same pipeline as used for *M. tuberculosis*. The raw SRA files (*n*=1,164) from the study by Grad et al. (Grad, 2014) were downloaded and aligned on the reference genome *N. gonorrhoeae* NCCP11945 (NC_011035.1) with BWA-MEM. The sequence of *N. lactamica* ATCC 23970 (SRR2906934) was used as a root for the phylogenetic analysis. Calculation of distances, bulding of the phylogenetic tree, parsimony and statistical analysis were performed as described above.

In the total set 64,574 single- and 707 (1.1%) dinucleotide substitutions were found. In total, 15 genes with statistically significant frequency of 2N mutations were identified (Table S3).

### Figure S1.

Phylogenetic tree of the 9941 genomes that were sequenced and analyzed by the CRYpTIC Consortium obtained by Nearest-Neighbour method rooted onto *M. canetii* (**A**). Main *M. tuberculosis* lineages are shown with different colors. Lineage 6 and animal-adapted strains (*M. tuberculosis var. bovis, var. orygis*) are tightly bound and comprise the most ancient branch. Sublineages were mapped onto the tree using the SNP lists from Coll, 2014 and Thawornwattana, 2021. The unadjusted distribution of isolates with substitutions are shown with color along the whole set of isolates.

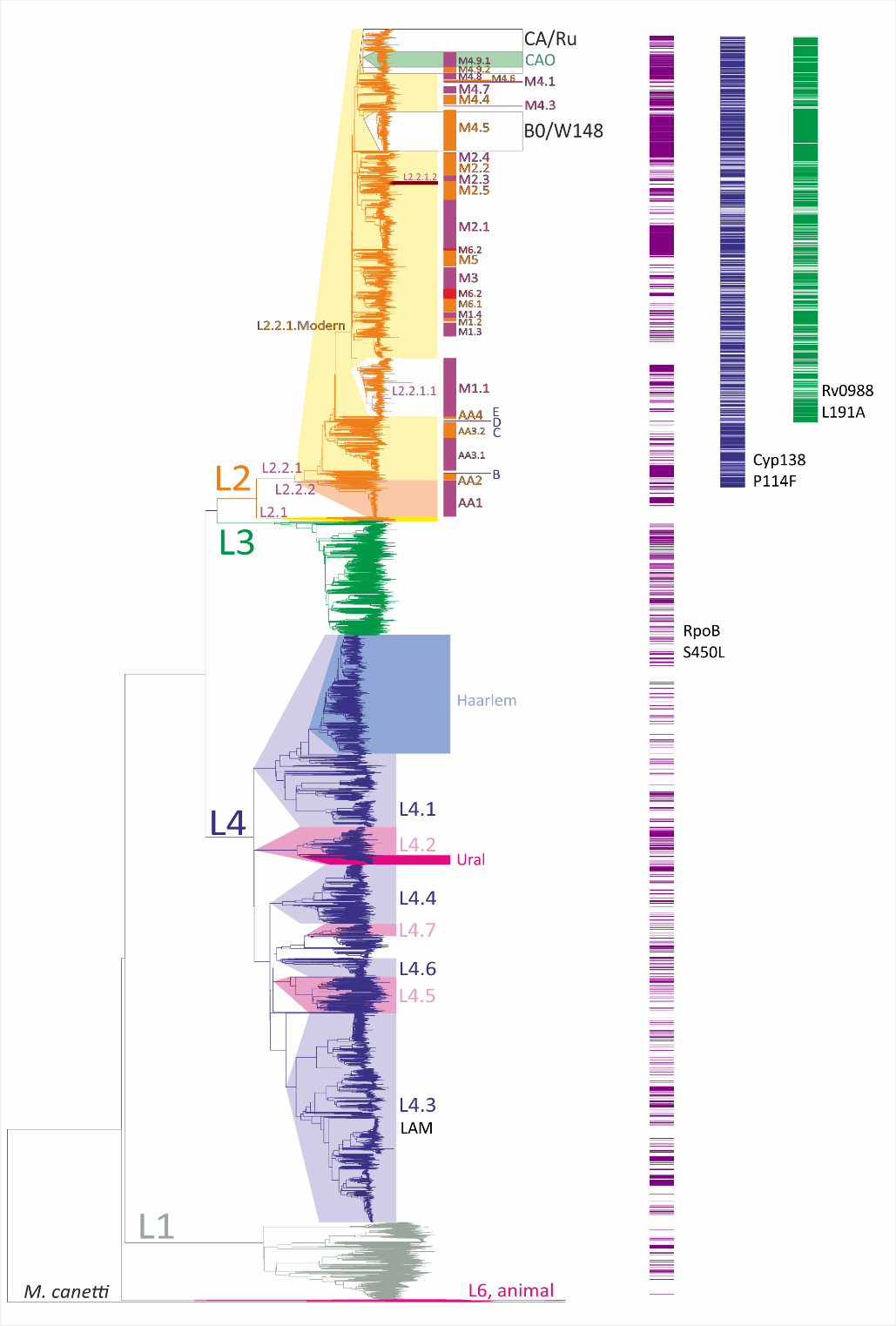

### Figure S2

Pairwise analysis of 2N vs. 1N substitution statistics for *M. tuberculosis* and *N. gonorrhoeae* sets of isolates. P-values for dN/dS are plotted on *x* axis, d2N/dS – on *y* axis. Statistical significance borderline with Bonferroni correction are shown as red dotted lines.

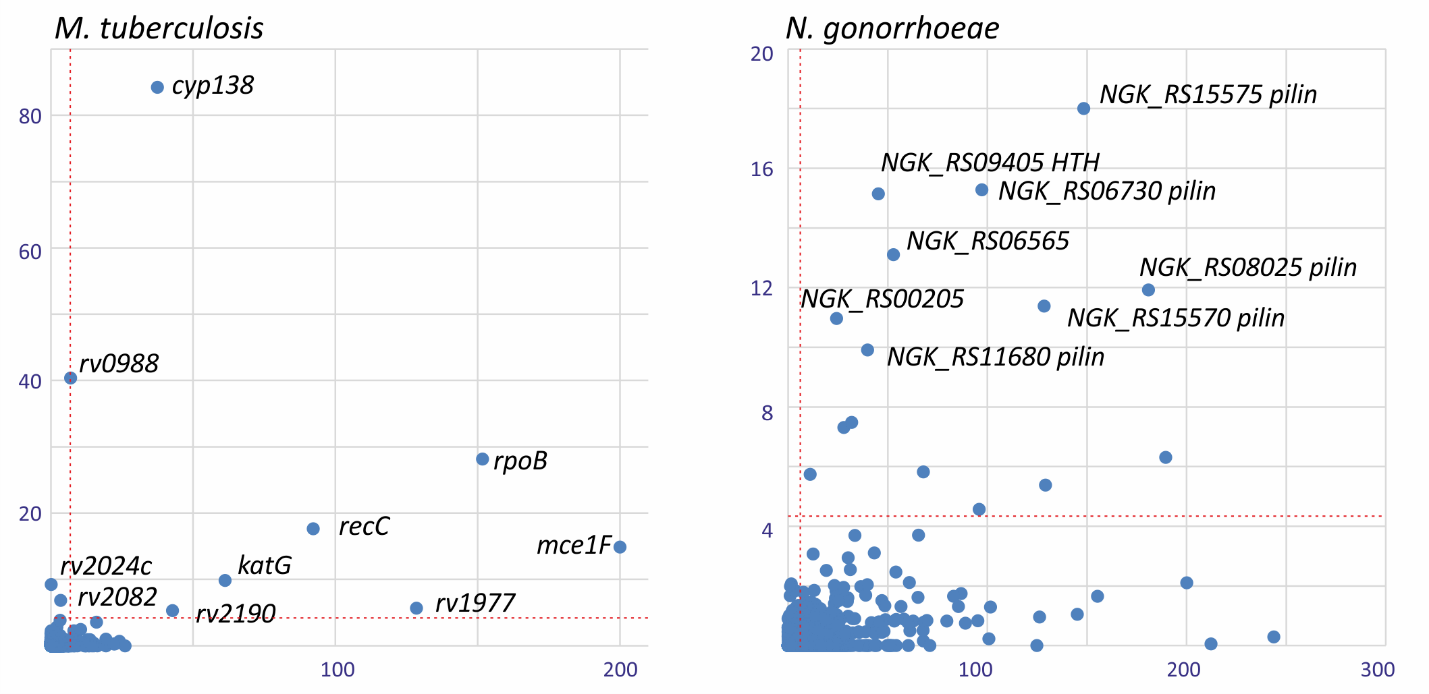

### Table S1.

RpoB substitutions and rifampicin resistance phenotype.

| Amino acid substitution | Wild-type codon | Mutated codon | Resistant isolates with mutation | Total number of resistan isolates in the study | Susceptible isolates with mutation | Total number of susceptible isolates in the study | Study |
| --- | --- | --- | --- | --- | --- | --- | --- |
| V170F | gtc | ttc | 35 | 71 |  |  | (Walker et al. 2022) |
| V170F | gtc | ttc | 3 | 3 | 0 | 3 | (Heep et al. 2000) |
| V170F | gtc | unkn. | 1 | 36 | 0 | 5 | (Jamieson et al. 2014) |
| V170F | gtc | unkn. | 5 | 871 | 0 | 6139 | (Zignol et al. 2018) |
| V170F | gtc | unkn. | 6 | 14 |  |  | (Solari, Santos-Lazaro, and Puyen 2020) |
| V170F | gtc | ttc | 2 | 324 | 0 | 28 | (Vargas et al. 2020) |
| V170G | gtc | ggc | 0 | 0 | 0 | 0 | (Walker et al. 2022) |
| V170A | gtc | gcc | 0 | 8 | 0 | 1 | (Walker et al. 2022) |
| Q172K | cag | aag | 0 | 1 | 0 | 0 | (Walker et al. 2022) |
| Q172R | cag | cgg | 0 | 7 | 0 | 0 | (Walker et al. 2022) |
| H194Y | cac | tac | 1 | 1 | 12 | 12 | (Walker et al. 2022) |
| H194R | cac | unkn. | 1 | 14 |  |  | (Solari, Santos-Lazaro, and Puyen 2020) |
| E250G | gag | ggg | 1 | 9 | 70 | 89 | (Walker et al. 2022) |
| E250G | gag | unkn. | 1 | 14 |  |  | (Solari, Santos-Lazaro, and Puyen 2020) |
| E250G | gag | ggg | 1 | 324 | 3 | 28 | (Vargas et al. 2020) |
| A286V | gcg | gtg | 0 | 41 | 0 | 0 | (Walker et al. 2022) |
| A286V | gcg | gtg | 1 | 21 | 0 | 19 | (Taniguchi et al. 1996) |
| Q409H | cag | cat | 1 | 41 |  |  | (Cavusoglu et al. 2002) |
| Q409R | cag | cgg | 0 | 38 | 0 | 0 | (Walker et al. 2022) |
| E423A | gag | gcg | 2 | 105 |  |  | (Schilke et al. 1999) |
| E423A | gag | gcg | 0 | 1 | 0 | 0 | (Walker et al. 2022) |
| E423G | gag | ggg | 0 | 1 | 0 | 0 | (Walker et al. 2022) |
| F424L | ttc | ctc | 0 | 4 | 0 | 0 | (Walker et al. 2022) |
| F424L | ttc | ttg | 2 | 13 | 0 | 26 | (Heym et al. 1994) |
| F424L | ttc | tta | 2 | 13 | 0 | 26 | (Stavrum et al. 2009) |
| F424L | ttc | ttg | 0 | 2 | 0 | 0 | (Walker et al. 2022) |
| F424S | ttc | tcc | 1 | 112 | 0 | 30 | (Wang et al. 2007) |
| F424V | ttc | gtc | 0 | 16 | 0 | 0 | (Walker et al. 2022) |
| F424C | ttc | tgc | 0 | 1 | 0 | 0 | (Walker et al. 2022) |
| G426S | ggc | agc | 1 | 41 | 0 | 245 | (Sandgren et al. 2009) |
| G426S | ggc | agc | 0 | 0 | 1 | 1 | (Walker et al. 2022) |
| G426D | ggc | gac | 1 | 32 | 0 | 26 | (Kim et al. 1997) |
| G426D | ggc | gat | 2 | 41 | 0 | 245 | (Sandgren et al. 2009) |
| G426G | ggc | ggt | 0 | 0 | 0 | 1 | (Walker et al. 2022) |
| T427G | acc | ggc | 0 | 0 | 1 | 1 | (Walker et al. 2022) |
| T427H | acc | cac | 6 | 41 | 0 | 245 | (Sandgren et al. 2009) |
| T427P | acc | ccc | 4 | 41 | 0 | 245 | (Sandgren et al. 2009) |
| T427A | acc | gcc | 2 | 41 | 0 | 245 | (Sandgren et al. 2009) |
| T427A | acc | gcc | 0 | 2 | 1 | 1 | (Walker et al. 2022) |
| T427I | acc | atc | 0 | 4 | 0 | 0 | (Walker et al. 2022) |
| T427N | acc | aac | 0 | 0 | 1 | 1 | (Walker et al. 2022) |
| T427S | acc | agc | 1 | 44 | 0 | 6 | (Sandgren et al. 2009) |
| T427S | acc | agc | 0 | 1 | 0 | 0 | (Walker et al. 2022) |
| T427S | acc | gct | 1 | 17 | 0 | 0 | (Sandgren et al. 2009) |
| T427T | acc | act | 0 | 0 | 0 | 2 | (Walker et al. 2022) |
| S428G | agc | ggc | 0 | 1 | 0 | 0 | (Walker et al. 2022) |
| S428Q | agc | cag | 1 | 15 | 0 | 10 | (Sandgren et al. 2009) |
| S428T | agc | acc | 0 | 3 | 0 | 0 | (Walker et al. 2022) |
| S428T | agc | acc | 1 | 108 | 0 | 12 | (Sandgren et al. 2009) |
| S428T | agc | unkn. | n.d. | 1302 | 1 |  | (Mvelase et al. 2019) |
| S428R | agc | agg | 0 | 28 | 1 | 110 | (Sandgren et al. 2009) |
| S428R | agc | cgc | 0 | 2 | 0 | 0 | (Walker et al. 2022) |
| S428R | agc | agg | 0 | 0 | 0 | 0 | (Walker et al. 2022) |
| S428R | agc | cgc | 1 | 57 | 0 |  | (ElMaraachli et al. 2015) |
| S428I | agc | atc | 0 | 1 | 0 | 0 | (Walker et al. 2022) |
| S428S | agc | agt | 0 | 1 | 0 | 0 | (Walker et al. 2022) |
| Q429H | cag | cac | 2 | 64 | 0 | 49 | (Sandgren et al. 2009) |
| Q429H | cag | cat | 0 | 2 | 0 | 0 | (Walker et al. 2022) |
| Q429H | cag | cac | 0 | 4 | 0 | 1 | (Walker et al. 2022) |
| Q429P | cag | unkn. | 1 | 36 | 0 | 0 | (Sirgel et al. 2013) |
| Q429P | cag | ccg | 0 | 0 | 0 | 0 | (Walker et al. 2022) |
| Q429K | cag | aag | 1 | 15 | 0 | 10 | (Sandgren et al. 2009) |
| Q429L | cag | ctg | 0 | 9 | 0 | 0 | (Walker et al. 2022) |
| L430P | ctg | unkn. | 2 | 66 | 0 | 56 | (Telenti et al. 1993) |
| L430P | ctg | ccg | 4 | 121 | 0 | 7 | (Sandgren et al. 2009) |
| L430P | ctg | ccg | 1 | 36 | 0 | 10 | (Bodmer et al. 1995) |
| L430P | ctg | ccg | 3 | 109 | 0 |  | (Wang et al. 2007) |
| L430P | ctg | unkn. | 1 | 300 | 2 | 315 | (Van Deun et al. 2013) |
| L430P | ctg | ccg | 0 | 36 | 2 | 5 | (Jamieson et al. 2014) |
| L430P | ctg | ccg | 0 | 102 | 3 | 62 | (Berrada et al. 2016) |
| L430P | ctg | ccg | 2 | 256 | 0 |  | (Jing et al. 2017) |
| L430P | ctg | unkn. | 3 | 139 | 1 | 17 | (Miotto et al. 2018) |
| L430P | ctg | unkn. | 10 | 871 | 20 | 6139 | (Zignol et al. 2018) |
| L430P | ctg | unkn. | n.d. | 1302 | 7 |  | (Mvelase et al. 2019) |
| L430P | ctg | ccg | 4 |  | 7 |  | (Torrea et al. 2019) |
| L430P | ctg | ccg | 31 | 106 | 96 | 103 | (Walker et al. 2022) |
| L430P | ctg | ccg | 0 | 153 | 1 | 9 | (Campbell et al. 2011) |
| L430R | ctg | cgg | 2 | 121 | 0 | 7 | (Sandgren et al. 2009) |
| L430R | ctg | cgg | 0 | 21 | 0 | 1 | (Walker et al. 2022) |
| L430V | ctg | gtg | 2 | 94 | 0 | 32 | (Sandgren et al. 2009) |
| L430V | ctg | unkn. | 0 | 300 | 1 | 315 | (Van Deun et al. 2013) |
| L430Q | ctg | unkn. | 1 | 139 | 0 | 17 | (Miotto et al. 2018) |
| S431T | agc | acc | 1 | 121 | 0 | 7 | (Sandgren et al. 2009) |
| S431T | agc | acc | 0 | 1 | 0 | 0 | (Walker et al. 2022) |
| S431R | agc | unkn. | 1 | 139 | 0 | 17 | (Miotto et al. 2018) |
| S431R | agc | cgc | 1 | 121 | 0 | 7 | (Sandgren et al. 2009) |
| S431R | agc | cgc | 0 | 2 | 0 | 0 | (Walker et al. 2022) |
| S431R | agc | aga | 0 | 0 | 0 | 0 | (Walker et al. 2022) |
| S431I | agc | atc | 1 | 64 | 0 | 49 | (Sandgren et al. 2009) |
| S431G | agc | ggc | 0 | 6 | 0 | 0 | (Walker et al. 2022) |
| Q432E | caa | gaa | 2 | 102 | 0 | 62 | (Berrada et al. 2016) |
| Q432E | caa | gaa | 1 | 154 | 0 | 113 | (Sandgren et al. 2009) |
| Q432E | caa | gaa | 1 | 139 | 0 | 17 | (Miotto et al. 2018) |
| Q432E | caa | gaa | 1 | 154 |  |  | (Matsui et al. 2020) |
| Q432E | caa | gaa | 0 | 1 | 0 | 0 | (Walker et al. 2022) |
| Q432H | caa | cac | 1 | 82 | 0 | 18 | (Sandgren et al. 2009) |
| Q432H | caa | cac | 0 | 2 | 0 | 0 | (Walker et al. 2022) |
| Q432H | caa | cat | 0 | 1 | 0 | 0 | (Walker et al. 2022) |
| Q432K | caa | aaa | 1 | 121 | 0 | 7 | (Sandgren et al. 2009) |
| Q432K | caa | aaa | 3 | 102 | 0 | 62 | (Berrada et al. 2016) |
| Q432K | caa | aaa | 1 | 49 | 0 | 114 | (Otchere et al. 2016) |
| Q432K | caa | aaa | 1 | 139 | 0 | 17 | (Miotto et al. 2018) |
| Q432K | caa | aaa | 4 | 336 | 0 | 55 | (Rodwell et al. 2014) |
| Q432K | caa | aaa | 4 | 57 |  |  | (ElMaraachli et al. 2015) |
| Q432K | caa | aaa | 2 | 871 | 1 | 6139 | (Zignol et al. 2018) |
| Q432K | caa | aaa | 18 | 34 | 1 | 1 | (Walker et al. 2022) |
| Q432L | caa | unkn. | 2 | 66 | 0 | 56 | (Telenti et al. 1993) |
| Q432L | caa | cta | 1 | 36 | 0 | 10 | (Bodmer et al. 1995) |
| Q432L | caa | cta | 1 | 10 | 0 |  | (Zaczek et al. 2009) |
| Q432L | caa | cta | 2 | 300 | 0 | 315 | (Van Deun et al. 2013) |
| Q432L | caa | cta | 2 | 121 | 0 | 7 | (Sandgren et al. 2009) |
| Q432L | caa | cta | 1 | 102 | 0 | 62 | (Berrada et al. 2016) |
| Q432L | caa | cta | 4 | 139 | 0 | 17 | (Miotto et al. 2018) |
| Q432L | caa | cta | 1 | 871 | 0 | 6139 | (Zignol et al. 2018) |
| Q432L | caa | unkn. | 1 | 52 |  |  | (Suthum, Samosornsuk, and Samosornsuk 2020) |
| Q432L | caa | cta | 16 | 21 | 1 | 1 | (Walker et al. 2022) |
| Q432L | caa | cta | 1 | 153 | 0 | 9 | (Campbell et al. 2011) |
| Q432N | caa | aat | 2 | 41 | 0 | 245 | (Sandgren et al. 2009) |
| Q432N | caa | aac | 1 | 256 | 0 |  | (Jing et al. 2017) |
| Q432N | caa | aat | 1 | 1 | 0 | 0 | (Walker et al. 2022) |
| Q432P | caa | cca | 1 | 41 |  |  | (Cavusoglu et al. 2002) |
| Q432P | caa | cca | 2 | 109 | 0 |  | (Wang et al. 2007) |
| Q432P | caa | cca | 2 | 300 | 0 | 315 | (Van Deun et al. 2013) |
| Q432P | caa | cca | 2 | 32 | 0 | 26 | (Sandgren et al. 2009) |
| Q432P | caa | cca | 2 | 102 | 0 | 62 | (Berrada et al. 2016) |
| Q432P | caa | cca | 1 | 49 | 0 | 114 | (Otchere et al. 2016) |
| Q432P | caa | cca | 6 | 139 | 0 | 17 | (Miotto et al. 2018) |
| Q432P | caa | cca | 2 | 871 | 0 | 6139 | (Zignol et al. 2018) |
| Q432P | caa | unkn. | n.d. | 1302 | 21 |  | (Mvelase et al. 2019) |
| Q432P | caa | cca | 1 | 324 | 0 | 28 | (Vargas et al. 2020) |
| Q432P | caa | cca | 20 | 29 | 0 | 1 | (Walker et al. 2022) |
| Q432Q | caa | unkn. | 1 | 139 | 0 | 17 | (Miotto et al. 2018) |
| Q432R | caa | cga |  |  |  |  | (Sandgren et al. 2009) |
| F433F | ttc | ttt | 0 | 102 | 2 | 62 | (Berrada et al. 2016) |
| F433F | ttc | ttt | 0 | 0 | 0 | 13 | (Walker et al. 2022) |
| F433L | ttc | ttg | 1 | 21 | 0 | 18 | (Sandgren et al. 2009) |
| F433V | ttc | gtc | 1 | 82 | 0 | 18 | (Sandgren et al. 2009) |
| M434I | atg | att | 1 | 3 | 1 | 0 | (Walker et al. 2022) |
| M434I | atg | ata |  |  |  |  | (Sandgren et al. 2009) |
| M434I | atg | ata | 1 | 8 | 1 | 2 | (Walker et al. 2022) |
| M434I | atg | atc | 1 | 5 | 1 | 1 | (Walker et al. 2022) |
| M434L | atg | ttg | 0 | 0 | 0 | 1 | (Walker et al. 2022) |
| M434V | atg | unkn. |  |  | 1 |  | (Yang et al. 1998) |
| M434V | atg | gtg | 0 | 4 | 0 | 3 | (Walker et al. 2022) |
| D435A | gac | gcc | 1 | 121 | 0 | 7 | (Sandgren et al. 2009) |
| D435A | gac | gcc | 1 | 84 | 0 | 36 | (Kambli et al. 2015) |
| D435A | gac | gcc | 0 | 6 | 1 | 2 | (Walker et al. 2022) |
| D435A | gac | gcc | 1 | 153 | 0 | 9 | (Campbell et al. 2011) |
| D435E | gac | gag | 1 | 163 |  |  | (Yang et al. 1998) |
| D435E | gac | gag | 2 | 121 | 0 | 7 | (Sandgren et al. 2009) |
| D435E | gac | gag | 0 | 2 | 0 | 0 | (Walker et al. 2022) |
| D435E | gac | gaa | 0 | 3 | 0 | 0 | (Walker et al. 2022) |
| D435F | gac | unkn. | 3 | 300 | 0 | 315 | (Van Deun et al. 2013) |
| D435F | gac | ttc | 1 | 84 | 0 | 36 | (Kambli et al. 2015) |
| D435F | gac | ttc | 1 | 102 | 0 | 62 | (Berrada et al. 2016) |
| D435F | gac | ttc | 4 | 871 | 0 | 6139 | (Zignol et al. 2018) |
| D435F | gac | ttc | 3 | 324 | 2 | 28 | (Vargas et al. 2020) |
| D435F | gac | ttc | 35 | 39 | 3 | 3 | (Walker et al. 2022) |
| D435G | gac | ggc | 3 | 109 | 0 |  | (Wang et al. 2007) |
| D435G | gac | ggc | 2 | 256 | 0 |  | (Jing et al. 2017) |
| D435G | gac | ggc | 2 | 871 | 0 | 6139 | (Zignol et al. 2018) |
| D435G | gac | ggc | 5 | 76 | 6 | 9 | (Walker et al. 2022) |
| D435H | gac | cac | 3 | 41 | 0 | 245 | (Sandgren et al. 2009) |
| D435H | gac | cac | 0 | 3 | 0 | 0 | (Walker et al. 2022) |
| D435I | gac | atc | 1 | 154 |  |  | (Matsui et al. 2020) |
| D435L | gac | unkn. | 1 | 139 | 0 | 17 | (Miotto et al. 2018) |
| D435L | gac | ctc | 2 | 2 | 0 | 0 | (Walker et al. 2022) |
| D435N | gac | aac | 1 | 105 | 0 | 0 | (Sandgren et al. 2009) |
| D435S | gac | unkn. | 4 | 36 |  |  | (Sirgel et al. 2013) |
| D435T | gac | unkn. | 1 | 36 |  |  | (Sirgel et al. 2013) |
| D435T | gac | acc | 4 | 69 |  |  | (Madania et al. 2012) |
| D435V | gac | unkn. | 6 | 66 | 0 | 56 | (Telenti et al. 1993) |
| D435V | gac | gtc | 1 | 121 | 0 | 7 | (Sandgren et al. 2009) |
| D435V | gac | gtc | 1 | 13 | 0 | 39 | (Heym et al. 1994) |
| D435V | gac | gtc | 3 | 36 | 0 | 10 | (Bodmer et al. 1995) |
| D435V | gac | gtc | 29 | 36 |  |  | (Sirgel et al. 2013) |
| D435V | gac | gtc | 18 | 102 | 0 | 62 | (Berrada et al. 2016) |
| D435V | gac | gtc | 1 | 10 | 0 |  | (Zaczek et al. 2009) |
| D435V | gac | gtc | 14 | 300 | 0 | 315 | (Van Deun et al. 2013) |
| D435V | gac | gtc | 3 | 84 | 0 | 36 | (Kambli et al. 2015) |
| D435V | gac | gtc | 3 |  | 0 |  | (Nosova et al. 2016) |
| D435V | gac | gtc | 1 | 139 | 0 | 17 | (Miotto et al. 2018) |
| D435V | gac | gtc | 2 | 163 |  |  | (Yang et al. 1998) |
| D435V | gac | gtc | 1 | 163 |  |  | (Yang et al. 1998) |
| D435V | gac | gtc | 1 | 163 |  |  | (Yang et al. 1998) |
| D435V | gac | gtc | 2 | 109 | 0 |  | (Wang et al. 2007) |
| D435V | gac | gtc | 23 | 336 | 0 | 55 | (Rodwell et al. 2014) |
| D435V | gac | gtc | 3 | 36 | 0 | 5 | (Jamieson et al. 2014) |
| D435V | gac | gtc | 15 | 57 |  |  | (ElMaraachli et al. 2015) |
| D435V | gac | gtc | 17 | 189 |  |  | (Rukasha et al. 2016) |
| D435V | gac | gtc | 11 | 49 | 0 | 114 | (Otchere et al. 2016) |
| D435V | gac | gtc | 5 | 256 | 0 |  | (Jing et al. 2017) |
| D435V | gac | unkn. | 30 | 871 | 5 | 6139 | (Zignol et al. 2018) |
| D435V | gac | unkn. | n.d. | 1302 | 16 |  | (Mvelase et al. 2019) |
| D435V | gac | unkn. | 1 | 52 |  |  | (Suthum, Samosornsuk, and Samosornsuk 2020) |
| D435V | gac | gtc | 1 | 154 |  |  | (Matsui et al. 2020) |
| D435V | gac | gtc | 67 | 324 | 3 | 28 | (Vargas et al. 2020) |
| D435V | gac | gtc | 706 | 732 | 8 | 9 | (Walker et al. 2022) |
| D435V | gac | gtc | 11 | 153 | 0 | 9 | (Campbell et al. 2011) |
| D435V | gac | gtc | 4 | 69 |  |  | (Madania et al. 2012) |
| D435Y | gac | tac | 2 | 36 | 0 | 10 | (Bodmer et al. 1995) |
| D435Y | gac | tac | 2 | 121 | 0 | 7 | (Sandgren et al. 2009) |
| D435Y | gac | tac | 1 | 109 | 0 |  | (Wang et al. 2007) |
| D435Y | gac | tac | 1 | 10 | 0 |  | (Zaczek et al. 2009) |
| D435Y | gac | tac | 7 | 300 | 1 | 315 | (Van Deun et al. 2013) |
| D435Y | gac | tac | 0 | 0 | 1 | 4 | (Williamson et al. 2012) |
| D435Y | gac | tac | 0 | 336 | 4 | 55 | (Rodwell et al. 2014) |
| D435Y | gac | tac | 3 | 84 | 0 | 36 | (Kambli et al. 2015) |
| D435Y | gac | tac | 0 | 57 | 2 |  | (ElMaraachli et al. 2015) |
| D435Y | gac | tac | 0 |  | 2 |  | (Nosova et al. 2016) |
| D435Y | gac | tac | 0 | 102 | 4 | 62 | (Berrada et al. 2016) |
| D435Y | gac | tac | 1 | 49 | 0 | 114 | (Otchere et al. 2016) |
| D435Y | gac | tac | 1 | 256 | 0 |  | (Jing et al. 2017) |
| D435Y | gac | tac | 5 | 139 | 3 | 17 | (Miotto et al. 2018) |
| D435Y | gac | tac | 12 | 871 | 12 | 6139 | (Zignol et al. 2018) |
| D435Y | gac | tac | 7 |  | 3 |  | (Torrea et al. 2019) |
| D435Y | gac | unkn. | n.d. | 1302 | 11 |  | (Mvelase et al. 2019) |
| D435Y | gac | tac | 2 | 154 |  |  | (Matsui et al. 2020) |
| D435Y | gac | tac | 63 | 162 | 37 | 44 | (Walker et al. 2022) |
| D435Y | gac | tac | 2 | 153 | 0 | 9 | (Campbell et al. 2011) |
| Q436Q | cag | caa | 2 | 57 |  |  | (ElMaraachli et al. 2015) |
| Q436N | cag | aac | 0 | 0 | 0 | 1 | (Walker et al. 2022) |
| Q436P | cag | ccg | 0 | 1 | 0 | 0 | (Walker et al. 2022) |
| N437D | aac | gac | 0 | 13 | 0 | 2 | (Walker et al. 2022) |
| N437H | aac | cac | 1 | 32 | 0 | 26 | (Sandgren et al. 2009) |
| N437H | aac | cac | 0 | 3 | 0 | 0 | (Walker et al. 2022) |
| N437I | aac | atc | 1 | 103 | 0 | 10 | (Sandgren et al. 2009) |
| N437I | aac | atc | 0 | 1 | 0 | 0 | (Walker et al. 2022) |
| N437S | aac | agc | 0 | 2 | 0 | 1 | (Walker et al. 2022) |
| N437T | aac | acc | 6 | 94 | 0 | 32 | (Sandgren et al. 2009) |
| N437Y | aac | tac | 1 | 112 | 0 | 0 | (Sandgren et al. 2009) |
| N437Y | aac | tac | 0 | 0 | 0 | 1 | (Walker et al. 2022) |
| N438K | aac | aag | 2 | 41 | 0 | 245 | (Sandgren et al. 2009) |
| L440L | ctg | ttg | 1 | 121 | 0 | 7 | (Sandgren et al. 2009) |
| L440L | ctg | ctt | 0 | 0 | 0 | 4 | (Walker et al. 2022) |
| L440M | ctg | atg | 1 | 121 | 0 | 7 | (Sandgren et al. 2009) |
| L440P | ctg | ccg | 0 | 21 | 1 | 18 | (Sandgren et al. 2009) |
| S441A | tcg | gcg | 0 | 1 | 0 | 0 | (Walker et al. 2022) |
| S441L | tcg | ttg | 1 | 6+6 | 0 | 56 | (Telenti et al. 1993) |
| S441L | tcg | unkn. | 1 | 36 | 0 | 10 | (Bodmer et al. 1995) |
| S441L | tcg | unkn. | 1 | 300 | 0 | 315 | (Van Deun et al. 2013) |
| S441L | tcg | ttg | 7 | 121 | 0 | 7 | (Sandgren et al. 2009) |
| S441L | tcg | ttg | 2 | 336 | 0 | 55 | (Rodwell et al. 2014) |
| S441L | tcg | ttg | 3 | 36 | 0 | 5 | (Jamieson et al. 2014) |
| S441L | tcg | ttg | 1 | 57 | 0 |  | (ElMaraachli et al. 2015) |
| S441L | tcg | ttg | 1 | 102 | 0 | 62 | (Berrada et al. 2016) |
| S441L | tcg | ttg | 1 | 49 | 0 | 114 | (Otchere et al. 2016) |
| S441L | tcg | ttg | 1 | 256 | 0 |  | (Jing et al. 2017) |
| S441L | tcg | ttg | 1 | 871 | 1 | 6139 | (Zignol et al. 2018) |
| S441L | tcg | ttg | 1 | 154 |  |  | (Matsui et al. 2020) |
| S441L | tcg | ttg | 16 | 26 | 0 | 0 | (Walker et al. 2022) |
| S441M | tcg | atg | 2 | 2 | 0 | 0 | (Walker et al. 2022) |
| S441Q | tcg | unkn. | 8 | 300 | 0 | 315 | (Van Deun et al. 2013) |
| S441Q | tcg | cag | 2 | 33 | 0 | 17 | (Sandgren et al. 2009) |
| S441Q | tcg | cag | 1 | 256 | 0 |  | (Jing et al. 2017) |
| S441Q | tcg | unkn. | 3 | 139 | 0 | 17 | (Miotto et al. 2018) |
| S441Q | tcg | unkn. | 1 | 871 | 0 | 6139 | (Zignol et al. 2018) |
| S441Q | tcg | cag | 11 | 11 | 0 | 0 | (Walker et al. 2022) |
| S441Q | tcg | cag | 2 | 153 | 0 | 9 | (Campbell et al. 2011) |
| S441V | tcg | gtg | 2 | 2 | 0 | 0 | (Walker et al. 2022) |
| S441W | tcg | tgg |  |  |  |  | (Sandgren et al. 2009) |
| S441W | tcg | unkn. | 2 | 139 | 0 | 17 | (Miotto et al. 2018) |
| S441W | tcg | tgg | 1 | 41 |  |  | (Cavusoglu et al. 2002) |
| S441W | tcg | tgg | 1 | 41 |  |  | (Cavusoglu et al. 2002) |
| S441P | tcg | ccg | 3 | 109 | 0 |  | (Wang et al. 2007) |
| G442A | ggg | gcg | 3 | 41 | 0 | 286 | (Bahrmand et al. 2009) |
| G442E | ggg | gag | 0 | 1 | 0 | 0 | (Walker et al. 2022) |
| G442W | ccc | acc | 1 | 101 | 0 | 21 | (Sandgren et al. 2009) |
| G442W | ggg | tgg | 1 | 37 | 0 | 0 | (Sandgren et al. 2009) |
| L443L | ttg | ttt | 0 | 2 | 1 | 1 | (Walker et al. 2022) |
| L443S | ttg | tcg | 1 | 15 | 0 | 10 | (Sandgren et al. 2009) |
| L443L | ttg | ctg | 0 | 0 | 0 | 1 | (Walker et al. 2022) |
| L443W | ttg | tgg | 1 | 82 | 0 | 18 | (Sandgren et al. 2009) |
| T444I | acc | atc | 1 | 37 | 0 | 0 | (Sandgren et al. 2009) |
| T444I | acc | atc | 0 | 2 | 0 | 0 | (Walker et al. 2022) |
| T444P | acc | ccc | 1 | 82 | 0 | 18 | (Sandgren et al. 2009) |
| T444S | acc | agc | 0 | 1 | 0 | 0 | (Walker et al. 2022) |
| T444T | acc | act | 0 | 0 | 0 | 1 | (Walker et al. 2022) |
| T444T | acc | acg | 0 | 4 | 0 | 0 | (Walker et al. 2022) |
| H445A | cac | gcc | 1 | 102 | 0 | 62 | (Berrada et al. 2016) |
| H445C | cac | tgc | 2 | 32 |  | 26 | (Sandgren et al. 2009) |
| H445C | cac | tgc | 0 | 41 | 1 |  | (Cavusoglu et al. 2002) |
| H445C | cac | tgc | 1 | 84 | 0 | 36 | (Kambli et al. 2015) |
| H445C | cac | tgc | 2 | 102 | 0 | 62 | (Berrada et al. 2016) |
| H445C | cac | tgc | 2 | 49 | 0 | 114 | (Otchere et al. 2016) |
| H445C | cac | tgc | 7 | 256 | 0 |  | (Jing et al. 2017) |
| H445C | cac | tgc | 3 | 139 | 0 | 17 | (Miotto et al. 2018) |
| H445C | cac | tgc | 1 | 871 | 0 | 6139 | (Zignol et al. 2018) |
| H445C | cac | tgc | 2 | 154 |  |  | (Matsui et al. 2020) |
| H445C | cac | tgc | 2 | 324 | 0 | 28 | (Vargas et al. 2020) |
| H445C | cac | tgc | 32 | 36 | 3 | 3 | (Walker et al. 2022) |
| H445C | cac | tgc | 1 | 69 |  |  | (Madania et al. 2012) |
| H445D | cac | unkn. | 5 | 66 | 0 | 56 | (Telenti et al. 1993) |
| H445D | cac | gac | 3 | 36 | 0 | 10 | (Bodmer et al. 1995) |
| H445D | cac | gac | 7 | 109 | 0 |  | (Wang et al. 2007) |
| H445D | cac | gac | 15 | 300 | 0 | 315 | (Van Deun et al. 2013) |
| H445D | cac | gac | 8 | 121 | 0 | 7 | (Sandgren et al. 2009) |
| H445D | cac | gac | 7 | 84 | 0 | 36 | (Kambli et al. 2015) |
| H445D | cac | gac | 1 | 10 | 0 |  | (Zaczek et al. 2009) |
| H445D | cac | gac | 1 |  | 0 |  | (Nosova et al. 2016) |
| H445D | cac | gac | 3 | 102 | 0 | 62 | (Berrada et al. 2016) |
| H445D | cac | gac | 3 | 139 | 0 | 17 | (Miotto et al. 2018) |
| H445D | cac | gac | 1 |  |  |  | (Springer et al. 2009) |
| H445D | cac | gac | 1 | 41 |  |  | (Cavusoglu et al. 2002) |
| H445D | cac | gac | 3 | 336 | 0 | 55 | (Rodwell et al. 2014) |
| H445D | cac | gac | 1 | 36 | 0 | 5 | (Jamieson et al. 2014) |
| H445D | cac | gac | 2 | 57 | 0 |  | (ElMaraachli et al. 2015) |
| H445D | cac | gac | 2 | 49 | 0 | 114 | (Otchere et al. 2016) |
| H445D | cac | gac | 19 | 189 |  |  | (Rukasha et al. 2016) |
| H445D | cac | gac | 11 | 256 | 0 |  | (Jing et al. 2017) |
| H445D | cac | gac | 1 | 256 | 0 |  | (Jing et al. 2017) |
| H445D | cac | gac | 46 | 871 | 4 | 6139 | (Zignol et al. 2018) |
| H445D | cac | unkn. | 1 | 52 |  |  | (Suthum, Samosornsuk, and Samosornsuk 2020) |
| H445D | cac | gac | 1 | 154 |  |  | (Matsui et al. 2020) |
| H445D | cac | gac | 7 | 324 | 0 | 28 | (Vargas et al. 2020) |
| H445D | cac | gac | 275 | 288 | 3 | 3 | (Walker et al. 2022) |
| H445D | cac | gac | 4 | 153 | 0 | 9 | (Campbell et al. 2011) |
| H445D | cac | gac | 4 | 69 |  |  | (Madania et al. 2012) |
| H445F | cac | ttc |  |  |  |  | (Bostanabad et al. 2007) |
| H445F | cac | ttc | 2 | 41 | 0 | 245 | (Sandgren et al. 2009) |
| H445F | cac | ttc | 1 | 871 | 0 | 6139 | (Zignol et al. 2018) |
| H445F | cac | ttc | 1 | 1 | 0 | 0 | (Walker et al. 2022) |
| H445G | cac | ggc | 3 | 121 | 0 | 7 | (Sandgren et al. 2009) |
| H445G | cac | ggc | 1 | 163 |  |  | (Yang et al. 1998) |
| H445G | cac | ggc | 1 | 102 | 0 | 62 | (Berrada et al. 2016) |
| H445G | cac | ggc | 4 | 7 | 3 | 3 | (Walker et al. 2022) |
| H445G | cac | ggc | 1 | 153 | 0 | 9 | (Campbell et al. 2011) |
| H445G | cac | ggc | 1 | 69 |  |  | (Madania et al. 2012) |
| H445L | cac | unkn. | 8 | 300 | 0 | 315 | (Van Deun et al. 2013) |
| H445L | cac | ctg | 1 | 82 | 0 | 18 | (Sandgren et al. 2009) |
| H445L | cac | unkn. | 18 | 871 | 4 | 6139 | (Zignol et al. 2018) |
| H445L | cac | ctc | 105 | 115 | 8 | 8 | (Walker et al. 2022) |
| H445L | cac | ctc | 2 | 121 | 0 | 7 | (Sandgren et al. 2009) |
| H445L | cac | ctc | 1 | 163 |  |  | (Yang et al. 1998) |
| H445L | cac | ctc | 11 | 109 | 0 |  | (Wang et al. 2007) |
| H445L | cac | ctc | 0 | 0 | 1 | 4 | (Williamson et al. 2012) |
| H445L | cac | ctc | 1 | 336 | 3 | 55 | (Rodwell et al. 2014) |
| H445L | cac | ctc | 1 | 36 | 1 | 5 | (Jamieson et al. 2014) |
| H445L | cac | ctc | 1 | 57 | 1 |  | (ElMaraachli et al. 2015) |
| H445L | cac | ctc | 1 |  | 1 |  | (Nosova et al. 2016) |
| H445L | cac | ctc | 1 | 189 |  |  | (Rukasha et al. 2016) |
| H445L | cac | ctc | 4 | 102 | 0 | 62 | (Berrada et al. 2016) |
| H445L | cac | ctc | 6 | 256 | 0 |  | (Jing et al. 2017) |
| H445L | cac | unkn. | n.d. | 1302 | 6 |  | (Mvelase et al. 2019) |
| H445L | cac | ctc | 5 | 154 |  |  | (Matsui et al. 2020) |
| H445L | cac | ctc | 2 | 324 | 0 | 28 | (Vargas et al. 2020) |
| H445L | cac | ctc | 3 | 153 | 2 | 9 | (Campbell et al. 2011) |
| H445L | cac | ctt | 1 | 49 | 0 | 7 | (Sandgren et al. 2009) |
| H445L | cac | unkn. | 7 | 139 | 2 | 17 | (Miotto et al. 2018) |
| H445N | cac | unkn. | 1 | 66 | 0 | 56 | (Telenti et al. 1993) |
| H445N | cac | aac | 3 | 109 | 0 |  | (Wang et al. 2007) |
| H445N | cac | aac | 3 | 46 | 0 | 286 | (Bahrmand et al. 2009) |
| H445N | cac | aac | 0 | 336 | 1 | 55 | (Rodwell et al. 2014) |
| H445N | cac | aac | 2 | 300 | 2 | 315 | (Van Deun et al. 2013) |
| H445N | cac | aac | 1 | 37 | 0 | 13 | (Sandgren et al. 2009) |
| H445N | cac | unkn. | 0 | 36 | 1 | 5 | (Jamieson et al. 2014) |
| H445N | cac | aac | 3 | 84 | 0 | 36 | (Kambli et al. 2015) |
| H445N | cac | aac | 0 |  | 1 |  | (Nosova et al. 2016) |
| H445N | cac | aac | 0 | 102 | 3 | 62 | (Berrada et al. 2016) |
| H445N | cac | aac | 4 | 256 | 0 |  | (Jing et al. 2017) |
| H445N | cac | unkn. | 4 | 139 | 3 | 17 | (Miotto et al. 2018) |
| H445N | cac | aac | 4 | 871 | 6 | 6139 | (Zignol et al. 2018) |
| H445N | cac | unkn. | 1 | 52 |  |  | (Suthum, Samosornsuk, and Samosornsuk 2020) |
| H445N | cac | aac | 1 | 154 |  |  | (Matsui et al. 2020) |
| H445N | cac | aac | 12 | 46 | 38 | 39 | (Walker et al. 2022) |
| H445N | cac | aac | 0 | 153 | 4 | 9 | (Campbell et al. 2011) |
| H445P | cac | unkn. | 1 | 66 | 0 | 56 | (Telenti et al. 1993) |
| H445P | cac | ccc | 2 | 26 |  | 10 | (Sandgren et al. 2009) |
| H445P | cac | ccc | 2 | 36 | 0 | 10 | (Bodmer et al. 1995) |
| H445P | cac | ccc | 1 | 189 |  |  | (Rukasha et al. 2016) |
| H445P | cac | ccc | 1 |  | 0 |  | (Nosova et al. 2016) |
| H445P | cac | unkn. | 2 | 139 | 0 | 17 | (Miotto et al. 2018) |
| H445P | cac | unkn. | 1 | 52 |  |  | (Suthum, Samosornsuk, and Samosornsuk 2020) |
| H445P | cac | ccc | 2 | 5 | 0 | 0 | (Walker et al. 2022) |
| H445Q | cac | cag | 2 | 26 | 0 | 10 | (Sandgren et al. 2009) |
| H445Q | cac | cag | 0 | 10 | 1 | 1 | (Walker et al. 2022) |
| H445Q | cac | caa | 1 | 121 | 0 | 7 | (Sandgren et al. 2009) |
| H445Q | cac | caa | 0 | 6 | 1 | 0 | (Walker et al. 2022) |
| H445Q | cac | unkn. | 5 | 139 | 0 | 17 | (Miotto et al. 2018) |
| H445R | cac | unkn. | 2 | 66 | 0 | 56 | (Telenti et al. 1993) |
| H445R | cac | cgc | 1 | 32 |  | 26 | (Sandgren et al. 2009) |
| H445R | cac | cgc | 2 | 36 | 0 | 10 | (Bodmer et al. 1995) |
| H445R | cac | cgc | 4 | 41 |  |  | (Cavusoglu et al. 2002) |
| H445R | cac | cgc | 4 | 109 | 0 |  | (Wang et al. 2007) |
| H445R | cac | cgc | 6 | 336 | 0 | 55 | (Rodwell et al. 2014) |
| H445R | cac | cgc | 5 | 300 | 0 | 315 | (Van Deun et al. 2013) |
| H445R | cac | cgc | 2 | 57 | 0 |  | (ElMaraachli et al. 2015) |
| H445R | cac | cgc | 1 | 84 | 0 | 36 | (Kambli et al. 2015) |
| H445R | cac | cgc | 2 | 102 | 0 | 62 | (Berrada et al. 2016) |
| H445R | cac | cgc | 3 |  | 0 |  | (Nosova et al. 2016) |
| H445R | cac | cgc | 2 | 49 | 0 | 114 | (Otchere et al. 2016) |
| H445R | cac | cgc | 1 | 189 |  |  | (Rukasha et al. 2016) |
| H445R | cac | cgc | 9 | 256 | 0 |  | (Jing et al. 2017) |
| H445R | cac | unkn. | 5 | 139 | 0 | 17 | (Miotto et al. 2018) |
| H445R | cac | cgc | 14 | 871 | 1 | 6139 | (Zignol et al. 2018) |
| H445R | cac | unkn. | n.d. | 1302 | 1 |  | (Mvelase et al. 2019) |
| H445R | cac | cgc | 1 | 154 |  |  | (Matsui et al. 2020) |
| H445R | cac | cgc | 64 | 79 | 2 | 2 | (Walker et al. 2022) |
| H445R | cac | cgc | 3 | 153 | 0 | 9 | (Campbell et al. 2011) |
| H445S | cac | agc | 0 | 102 | 2 | 62 | (Berrada et al. 2016) |
| H445S | cac | tcc | 0 | 102 | 1 | 62 | (Berrada et al. 2016) |
| H445S | cac | agc | 1 | 154 |  |  | (Sandgren et al. 2009) |
| H445S | cac | unkn. | 0 | 871 | 2 | 6139 | (Zignol et al. 2018) |
| H445S | cac | unkn. | 3 | 139 | 4 | 17 | (Miotto et al. 2018) |
| H445S | cac | unkn. | n.d. | 1302 | 2 |  | (Mvelase et al. 2019) |
| H425S | cac | tcc | 6 | 14 | 2 | 4 | (Walker et al. 2022) |
| H425S | cac | agc | 2 | 14 | 0 | 2 | (Walker et al. 2022) |
| H445T | cac | acc | 1 | 121 |  | 7 | (Sandgren et al. 2009) |
| H445T | cac | acc | 1 | 871 | 0 | 6139 | (Zignol et al. 2018) |
| H425T | cac | acc | 3 | 3 | 0 | 0 | (Walker et al. 2022) |
| H445Y | cac | unkn. | 8 | 66 | 0 | 56 | (Telenti et al. 1993) |
| H445Y | cac | tac | 39 | 121 | 0 | 7 | (Sandgren et al. 2009) |
| H445Y | cac | tac | 3 | 36 | 0 | 10 | (Bodmer et al. 1995) |
| H445Y | cac | tac | 1 | 163 |  |  | (Yang et al. 1998) |
| H445Y | cac | tac | 2 | 41 |  |  | (Cavusoglu et al. 2002) |
| H445Y | cac | tac | 9 | 109 | 0 |  | (Wang et al. 2007) |
| H445Y | cac | tac | 3 | 41 | 0 | 286 | (Bahrmand et al. 2009) |
| H445Y | cac | tac | 18 | 336 | 0 | 55 | (Rodwell et al. 2014) |
| H445Y | cac | tac | 2 | 36 | 0 | 5 | (Jamieson et al. 2014) |
| H445Y | cac | tac | 20 | 300 | 1 | 315 | (Van Deun et al. 2013) |
| H445Y | cac | tac | 3 | 84 | 0 | 36 | (Kambli et al. 2015) |
| H445Y | cac | tac | 1 | 49 | 0 | 114 | (Otchere et al. 2016) |
| H445Y | cac | tac | 3 | 57 | 0 |  | (ElMaraachli et al. 2015) |
| H445Y | cac | tac | 27 | 189 |  |  | (Rukasha et al. 2016) |
| H445Y | cac | tac | 2 |  | 0 |  | (Nosova et al. 2016) |
| H445Y | cac | tac | 1 | 102 | 0 | 62 | (Berrada et al. 2016) |
| H445Y | cac | tac | 2 | 256 | 0 |  | (Jing et al. 2017) |
| H445Y | cac | tac | 31 | 871 | 3 | 6139 | (Zignol et al. 2018) |
| H445Y | cac | unkn. | 1 | 139 | 0 | 17 | (Miotto et al. 2018) |
| H445Y | cac | unkn. | 4 | 52 |  |  | (Suthum, Samosornsuk, and Samosornsuk 2020) |
| H445Y | cac | tac | 4 | 154 |  |  | (Matsui et al. 2020) |
| H445Y | cac | tac | 6 | 324 | 0 | 28 | (Vargas et al. 2020) |
| H445Y | cac | tac | 305 | 347 | 2 | 4 | (Walker et al. 2022) |
| H445Y | cac | tac | 18 | 153 | 0 | 9 | (Campbell et al. 2011) |
| H445Y | cac | tac | 2 | 69 |  |  | (Madania et al. 2012) |
| K446E | aag | gag | 0 | 1 | 0 | 0 | (Walker et al. 2022) |
| K446Q | aag | cag | 1 | 108 | 0 | 12 | (Sandgren et al. 2009) |
| K446Q | aag | cag | 0 | 7 | 0 | 0 | (Walker et al. 2022) |
| K446N | aag | aat | 2 | 94 | 0 | 32 | (Sandgren et al. 2009) |
| K446R | aag | agg | 0 | 2 | 0 | 0 | (Walker et al. 2022) |
| K446T | aag | acg | 0 | 1 | 0 | 0 | (Walker et al. 2022) |
| R447H | cgc | cac | 2 | 94 | 0 | 32 | (Sandgren et al. 2009) |
| R447P | cgc | ccc | 4 | 94 | 0 | 32 | (Sandgren et al. 2009) |
| R447P | cgc | cct | 1 | 15 |  | 10 | (Sandgren et al. 2009) |
| R447R | cgc | cgt | 0 | 0 | 0 | 53 | (Walker et al. 2022) |
| R448Q | cga | caa | 2 | 6 | 0 | 0 | (Walker et al. 2022) |
| R448K | cga | unkn. | 1 | 871 | 0 | 6139 | (Zignol et al. 2018) |
| R448K | cga | aaa | 2 | 2 | 0 | 0 | (Walker et al. 2022) |
| R448R | cga | cgg | 0 | 0 | 0 | 1 | (Walker et al. 2022) |
| L449L | ctg | ctc | 0 | 2 | 0 | 0 | (Walker et al. 2022) |
| L449L | ctg | cta | 0 | 0 | 0 | 2 | (Walker et al. 2022) |
| L449M | ctg | atg | 0 | 9 | 0 | 0 | (Walker et al. 2022) |
| L449P | ctg | unkn. | 2 |  |  |  | (Singh et al. 2020) |
| S450C | tcg | gcg | 0 | 1 | 0 | 0 | (Walker et al. 2022) |
| S450C | tcg | tgt | 0 | 102 | 1 | 62 | (Berrada et al. 2016) |
| S450C | tcg | tgt | 1 | 121 | 0 | 7 | (Sandgren et al. 2009) |
| S450C | tcg | unkn. | 1 | 871 | 1 | 6139 | (Zignol et al. 2018) |
| S450C | tcg | tgt | 0 | 0 | 1 | 1 | (Walker et al. 2022) |
| S450C | tcg | tgc | 0 | 1 | 0 | 0 | (Walker et al. 2022) |
| S450F | tcg | ttc | 1 | 102 | 0 | 62 | (Berrada et al. 2016) |
| S450F | tcg | ttt | 2 | 103 | 0 | 10 | (Sandgren et al. 2009) |
| S450F | tcg | unkn. | 2 | 139 | 0 | 17 | (Miotto et al. 2018) |
| S450F | tcg | ttc | 38 | 871 | 0 | 6139 | (Zignol et al. 2018) |
| S450F | tcg | ttc | 6 | 324 | 0 | 28 | (Vargas et al. 2020) |
| S450F | tcg | ttc | 70 | 77 | 0 | 0 | (Walker et al. 2022) |
| S450F | tcg | ttc | 1 | 153 | 0 | 9 | (Campbell et al. 2011) |
| S450F | tcg | ttt | 33 | 35 | 0 | 0 | (Walker et al. 2022) |
| S450G | tcg | ggg | 0 | 1 | 0 | 0 | (Walker et al. 2022) |
| S450G | tcg | ggg | 1 | 69 |  |  | (Madania et al. 2012) |
| S450L | tcg | unkn. | 31 | 66 | 0 | 56 | (Telenti et al. 1993) |
| S450L | tcg | ttg | 1 | 1 | 0 | 8 | (Sandgren et al. 2009) |
| S450L | tcg | ttg | 4 | 13 | 0 | 39 | (Heym et al. 1994) |
| S450L | tcg | ttg | 3 | 36 | 0 | 10 | (Bodmer et al. 1995) |
| S450L | tcg | ttg | 33 | 109 | 0 |  | (Wang et al. 2007) |
| S450L | tcg | ttg | 1 | 10 | 0 |  | (Zaczek et al. 2009) |
| S450L | tcg | ttg | 14 | 163 |  |  | (Yang et al. 1998) |
| S450L | tcg | ttg | 1 | 163 |  |  | (Yang et al. 1998) |
| S450L | tcg | ttg | 41 | 163 |  |  | (Yang et al. 1998) |
| S450L | tcg | ttg | 15 | 41 |  |  | (Cavusoglu et al. 2002) |
| S450L | tcg | ttg | 1 | 41 |  |  | (Cavusoglu et al. 2002) |
| S450L | tcg | ttg | 1 | 41 |  |  | (Cavusoglu et al. 2002) |
| S450L | tcg | ttg | 1 | 41 |  |  | (Cavusoglu et al. 2002) |
| S450L | tcg | ttg | 1 | 41 |  |  | (Cavusoglu et al. 2002) |
| S450L | tcg | ttg | 10 |  |  |  | (Springer et al. 2009) |
| S450L | tcg | ttg | 1 | 36 |  |  | (Sirgel et al. 2013) |
| S450L | tcg | ttg | 231 | 336 | 1 | 55 | (Rodwell et al. 2014) |
| S450L | tcg | unkn. | 170 | 300 | 9 | 315 | (Van Deun et al. 2013) |
| S450L | tcg | ttg | 13 |  |  |  | (Htike Min et al. 2014) |
| S450L | tcg | ttg | 19 | 36 | 0 | 5 | (Jamieson et al. 2014) |
| S450L | tcg | ttg | 14 | 57 | 0 |  | (ElMaraachli et al. 2015) |
| S450L | tcg | ttg | 26 | 49 | 0 | 114 | (Otchere et al. 2016) |
| S450L | tcg | ttg | 57 | 84 | 0 | 36 | (Kambli et al. 2015) |
| S450L | tcg | ttg | 93 |  | 0 |  | (Nosova et al. 2016) |
| S450L | tcg | ttg | 105 | 189 |  |  | (Rukasha et al. 2016) |
| S450L | tcg | ttg | 5 | 102 | 0 | 62 | (Berrada et al. 2016) |
| S450L | tcg | ttg | 156 | 256 | 0 |  | (Jing et al. 2017) |
| S450L | tcg | unkn. | 4 | 139 | 0 | 17 | (Miotto et al. 2018) |
| S450L | tcg | unkn. | 530 | 871 | 11 | 6139 | (Zignol et al. 2018) |
| S450L | tcg | ttg | 5 |  | 0 |  | (Torrea et al. 2019) |
| S450L | tcg | ttg | n.d. | 1302 | 3 |  | (Mvelase et al. 2019) |
| S450L | tcg | unkn. | 41 | 52 |  |  | (Suthum, Samosornsuk, and Samosornsuk 2020) |
| S450L | tcg | ttg | 107 | 154 |  |  | (Matsui et al. 2020) |
| S450L | tcg | ttg | 120 | 324 | 2 | 28 | (Vargas et al. 2020) |
| S450L | tcg | ttg | 5332 | 6536 | 67 | 74 | (Walker et al. 2022) |
| S450L | tcg | ttg | 101 | 153 | 0 | 9 | (Campbell et al. 2011) |
| S450L | tcg | ttg | 39 | 69 |  |  | (Madania et al. 2012) |
| S450L | tcg | ctg | 1 | 1 | 0 | 0 | (Walker et al. 2022) |
| S450M | tcg | atg | 1 | 324 | 2 | 28 | (Vargas et al. 2020) |
| S450M | tcg | atg | 5 | 5 | 0 | 0 | (Walker et al. 2022) |
| S450P | tcg | unkn. | n.d. | 1302 | 1 |  | (Mvelase et al. 2019) |
| S450Q | tcg | unkn. | 1 | 66 | 0 | 56 | (Telenti et al. 1993) |
| S450Q | tcg | cag | 1 | 121 | 0 | 7 | (Sandgren et al. 2009) |
| S450Q | tcg | cag | 1 | 189 |  |  | (Rukasha et al. 2016) |
| S450Q | tcg | cag | 1 | 256 | 0 |  | (Jing et al. 2017) |
| S450Q | tcg | unkn. | 1 | 139 | 0 | 17 | (Miotto et al. 2018) |
| S450Q | tcg | cag | 6 | 8 | 0 | 0 | (Walker et al. 2022) |
| S450V | tcg | gtg | 0 | 2 | 1 | 1 | (Walker et al. 2022) |
| S450W | tcg | unkn. | 1 | 66 | 0 | 56 | (Telenti et al. 1993) |
| S450W | tcg | tgg | 1 | 26 | 0 | 10 | (Bodmer et al. 1995) |
| S450W | tcg | tgg | 15 | 109 | 0 |  | (Wang et al. 2007) |
| S450W | tcg | tgg | 4 | 41 |  |  | (Cavusoglu et al. 2002) |
| S450W | tcg | unkn. | 3 | 300 | 0 | 315 | (Van Deun et al. 2013) |
| S450W | tcg | tgg | 7 | 336 | 0 | 55 | (Rodwell et al. 2014) |
| S450W | tcg | tgg | 3 | 36 | 0 | 5 | (Jamieson et al. 2014) |
| S450W | tcg | tgg | 5 | 57 | 0 |  | (ElMaraachli et al. 2015) |
| S450W | tcg | tgg | 1 |  | 0 |  | (Nosova et al. 2016) |
| S450W | tcg | tgg | 3 | 102 | 0 | 62 | (Berrada et al. 2016) |
| S450W | tcg | unkn. | 22 | 139 | 0 | 17 | (Miotto et al. 2018) |
| S450W | tcg | tgg | 9 | 871 | 0 | 6139 | (Zignol et al. 2018) |
| S450W | tcg | unkn. | 1 | 52 |  |  | (Suthum, Samosornsuk, and Samosornsuk 2020) |
| S450W | tcg | tgg | 8 | 154 |  |  | (Matsui et al. 2020) |
| S450W | tcg | tgg | 2 | 324 | 0 | 28 | (Vargas et al. 2020) |
| S450W | tcg | tgg | 127 | 151 | 5 | 5 | (Walker et al. 2022) |
| S450W | tcg | tgg | 2 | 153 | 0 | 9 | (Campbell et al. 2011) |
| S450W | tcg | tgg | 2 | 69 |  |  | (Madania et al. 2012) |
| S450Y | tcg | unkn. | 1 | 36 | 0 | 10 | (Bodmer et al. 1995) |
| S450Y | tcg | unkn. | 1 | 139 | 0 | 17 | (Miotto et al. 2018) |
| S450Y | tcg | tac | 2 | 2 | 0 | 0 | (Walker et al. 2022) |
| A451G | gcg | ggg | 0 | 1 | 0 | 0 | (Walker et al. 2022) |
| A451G | gcg | ggc | 0 | 1 | 0 | 0 | (Walker et al. 2022) |
| A451V | gcg | gtg | 0 | 7 | 4 | 4 | (Walker et al. 2022) |
| L452Q | ctg | cag | 0 | 0 | 1 | 1 | (Walker et al. 2022) |
| L452L | ctg | ttg | 0 | 0 | 0 | 1 | (Walker et al. 2022) |
| L452L | ctg | ctt | 0 | 1 | 0 | 1 | (Walker et al. 2022) |
| L452M | ctg | atg | 0 | 0 | 1 | 1 | (Walker et al. 2022) |
| L452P | ctg | unkn. | 1 | 66 | 0 | 56 | (Telenti et al. 1993) |
| L452P | ctg | ccg | 1 | 24 | 0 | 0 | (Sandgren et al. 2009) |
| L452P | ctg | ccg |  |  |  |  | (Yang et al. 1998) |
| L452P | ctg | ccg | 0 | 41 | 1 |  | (Cavusoglu et al. 2002) |
| L452P | ctg | ccg | 0 | 41 | 1 |  | (Cavusoglu et al. 2002) |
| L452P | ctg | unkn. | 9 | 300 | 1 | 315 | (Van Deun et al. 2013) |
| L452P | ctg | ccg |  |  |  |  | (Somoskovi et al. 2013) |
| L452P | ctg | ccg | 4 | 84 | 0 | 36 | (Kambli et al. 2015) |
| L452P | ctg | ccg | 0 | 102 | 2 | 62 | (Berrada et al. 2016) |
| L452P | ctg | ccg | 3 | 256 | 0 |  | (Jing et al. 2017) |
| L452P | ctg | unkn. | 11 | 139 | 2 | 17 | (Miotto et al. 2018) |
| L452P | ctg | unkn. | 12 | 871 | 15 | 6139 | (Zignol et al. 2018) |
| L452P | ctg | ccg | 6 |  | 3 |  | (Torrea et al. 2019) |
| L452P | ctg | unkn. | 1 | 14 |  |  | (Solari, Santos-Lazaro, and Puyen 2020) |
| L452P | ctg | unkn. | 1 | 52 |  |  | (Suthum, Samosornsuk, and Samosornsuk 2020) |
| L452P | ctg | ccg | 1 | 154 |  |  | (Matsui et al. 2020) |
| L452P | ctg | unkn. | 0 | 324 | 4 | 28 | (Vargas et al. 2020) |
| L452P | ctg | ccg | 78 | 121 | 50 | 53 | (Walker et al. 2022) |
| L452P | ctg | ccg | 1 | 153 | 2 | 9 | (Campbell et al. 2011) |
| L452P | ctg | ccg | 3 | 69 |  |  | (Madania et al. 2012) |
| L452P | ctg | cct | 2 | 82 | 0 | 18 | (Sandgren et al. 2009) |
| L452P | ctg | ccc | 4 | 189 |  |  | (Rukasha et al. 2016) |
| L452V | ctg | gtg | 0 | 1 | 0 | 0 | (Walker et al. 2022) |
| P454H | ccc | cac | 1 | 154 | 0 | 113 | (Sandgren et al. 2009) |
| P454H | ccc | cac | 0 | 2 | 2 | 2 | (Walker et al. 2022) |
| P454L | ccc | ctc | 0 | 0 | 6 | 6 | (Walker et al. 2022) |
| P454R | ccc | cgc | 0 | 1 | 0 | 0 | (Walker et al. 2022) |
| P454S |  | unkn. | 0 | 300 | 3 | 315 | (Van Deun et al. 2013) |
| P454S | ccc | tcc | 4 | 51 | 0 | 19 | (Sandgren et al. 2009) |
| P454S | ccc | tcc | 0 | 1 | 1 | 1 | (Walker et al. 2022) |
| G455G | ggc | ggg | 4 |  |  |  | (Htike Min et al. 2014) |
| G455G | ggc | ggt | 0 | 0 | 0 | 2 | (Walker et al. 2022) |
| L457R | ctg | cgt | 1 |  |  |  | (Htike Min et al. 2014) |
| L457P | ctg | ccg | 1 | 112 | 0 | 30 | (Sandgren et al. 2009) |
| E460G | gag | ggg | 1 | 154 | 0 | 113 | (Sandgren et al. 2009) |
| E460G | gag | gat | 1 | 37 |  |  | (Sandgren et al. 2009) |
| E460G | gag | ggg | 0 | 5 | 0 | 2 | (Walker et al. 2022) |
| L464L | ctg | ttg | 0 | 0 | 0 | 1 | (Walker et al. 2022) |
| L464M | ctc | atg | 3 | 19 | 0 | 22 | (Sandgren et al. 2009) |
| S472A | tcg | gcg | 1 | 37 | 0 | 0 | (Sandgren et al. 2009) |
| I480I | atc | att | 0 | 0 | 0 | 1 | (Walker et al. 2022) |
| I480T | atc | acc | 1 | 111 | 0 | 0 | (Sandgren et al. 2009) |
| I480T | atc | acc | 0 | 3 | 0 | 0 | (Walker et al. 2022) |
| I480V | atc | gtc | 1 | 37 | 0 | 13 | (Sandgren et al. 2009) |
| I480V | atc | gtc | 0 | 43 | 0 | 2 | (Walker et al. 2022) |
| E481A | gaa | gca | 0 | 8 | 0 | 0 | (Walker et al. 2022) |
| E481E | gaa | gag | 0 | 0 | 0 | 1 | (Walker et al. 2022) |
| E481G | gaa | gga | 1 | 122 |  |  | (Sandgren et al. 2009) |
| P483S | cct | tct | 0 | 2 | 0 | 0 | (Walker et al. 2022) |
| P483L | cct | ctt | 1 | 122 |  |  | (Sandgren et al. 2009) |
| P483L | cct | ctt | 0 | 2 | 1 | 1 | (Walker et al. 2022) |
| N487H | acc | cac | 0 | 2 | 0 | 0 | (Walker et al. 2022) |
| N487S | acc | agc | 0 | 3 | 0 | 1 | (Walker et al. 2022) |
| N487N | acc | aat | 0 | 4 | 0 | 1 | (Walker et al. 2022) |
| L490L | ctg | ttg | 0 | 0 | 0 | 1 | (Walker et al. 2022) |
| L490L | ctg | ctt | 0 | 0 | 0 | 3 | (Walker et al. 2022) |
| L490V | ctg | gtg | 2 | 111 |  |  | (Sandgren et al. 2009) |
| L490V | ctg | gtg | 0 | 2 | 0 | 0 | (Walker et al. 2022) |
| I491F | atc | unkn. | 5 | 300 | 0 | 315 | (Van Deun et al. 2013) |
| I491F | atc | ttc | 1 | 33 |  | 17 | (Sandgren et al. 2009) |
| I491F | atc | ttc | 3 | 256 | 0 |  | (Jing et al. 2017) |
| I491F | atc | unkn. | 3 | 871 | 2 | 6139 | (Zignol et al. 2018) |
| I491F | atc | unkn. | 6 |  | 2 |  | (Torrea et al. 2019) |
| I491F | atc | unkn. | 2 |  |  |  | (Solari, Santos-Lazaro, and Puyen 2020) |
| I491F | atc | ttc | 1 | 324 | 2 | 28 | (Vargas et al. 2020) |
| I491F | atc | unkn. |  |  |  |  | (Ardizzoni et al. 2021) |
| I491F | atc | ttc | 45 | 54 | 53 | 57 | (Walker et al. 2022) |
| I491F | atc | ttc | 1 | 69 |  |  | (Madania et al. 2012) |
| I491L | atc | ctc | 0 | 25 | 0 | 0 | (Walker et al. 2022) |
| I491M | atc | atg | 2 | 14 | 2 | 3 | (Walker et al. 2022) |
| I491S | atc | agc | 0 | 1 | 0 | 0 | (Walker et al. 2022) |
| I491T | atc | acc | 0 | 6 | 0 | 0 | (Walker et al. 2022) |
| I491V | atc | gtc | 0 | 8 | 0 | 0 | (Walker et al. 2022) |
| I491Y | atc | tac | 1 | 154 |  |  | (Matsui et al. 2020) |
| I491Y | atc | tac | 3 | 3 | 0 | 0 | (Walker et al. 2022) |
| S493L | tcg | ttg | 1 | 221 | 0 | 29 | (Sandgren et al. 2009) |
| S493L | tcg | ttg | 0 | 5 | 0 | 1 | (Walker et al. 2022) |
| S493W | tcg | tgg | 0 | 0 | 3 | 3 | (Walker et al. 2022) |
| L494L | ctg | cta | 0 | 1 | 0 | 11 | (Walker et al. 2022) |
| S541A | tcg | gcg | 1 | 154 | 0 | 113 | (Sandgren et al. 2009) |
| R552C | cgc | tgc | 1 | 111 |  |  | (Sandgren et al. 2009) |
| G591D | gag | gat | 1 | 111 |  |  | (Sandgren et al. 2009) |
| A692T | gcc | acc | 1 | 324 | 0 | 28 | (Vargas et al. 2020) |
| V695L | gtg | unkn. | 1 | 14 |  |  | (Solari, Santos-Lazaro, and Puyen 2020) |
| V695L | gtg | ctg | 1 | 324 | 4 | 28 | (Vargas et al. 2020) |
| R741S | cac | tgt | 1 | 324 | 0 | 28 | (Vargas et al. 2020) |
| I965V | unkn. | unkn. | 0 | 324 | 3 | 28 | (Vargas et al. 2020) |
| Q980R | unkn. | unkn. | 1 | 324 | 0 | 28 | (Vargas et al. 2020) |
| R1163H | cgc | cac | 0 | 324 | 1 | 28 | (Vargas et al. 2020) |

Ardizzoni, E., E. Ariza, D. Mulengwa, Q. Mpala, R. de La Tour, G. Maphalala, F. Varaine, et al. 2021. “Thin Layer Agar-Based Direct Phenotypic Drug-Susceptibility Testing on Sputum in Eswatini Rapidly Detects Mycobacterium Tuberculosis Growth and Rifampicin Resistance, Otherwise Missed by WHO Endorsed Diagnostic Tests.” *Antimicrobial Agents and Chemotherapy*, March. https://doi.org/10.1128/AAC.02263-20.

Bahrmand, Ahmad Reza, Leonid P. Titov, Alireza Hadizadeh Tasbiti, Shamsi Yari, and Edward A. Graviss. 2009. “High-Level Rifampin Resistance Correlates with Multiple Mutations in the rpoB Gene of Pulmonary Tuberculosis Isolates from the Afghanistan Border of Iran.” *Journal of Clinical Microbiology* 47 (9): 2744–50. https://doi.org/10.1128/JCM.r00548-09.

Berrada, Zenda L., Shou-Yean Grace Lin, Timothy C. Rodwell, Duylinh Nguyen, Gisela F. Schecter, Lucy Pham, J. Michael Janda, Wael Elmaraachli, Antonino Catanzaro, and Edward Desmond. 2016. “Rifabutin and Rifampin Resistance Levels and Associated rpoB Mutations in Clinical Isolates of Mycobacterium Tuberculosis Complex.” *Diagnostic Microbiology and Infectious Disease* 85 (2): 177–81. https://doi.org/10.1016/j.diagmicrobio.2016.01.019.

Bodmer, T., G. Zürcher, P. Imboden, and A. Telenti. 1995. “Mutation Position and Type of Substitution in the Beta-Subunit of the RNA Polymerase Influence in-Vitro Activity of Rifamycins in Rifampicin-Resistant Mycobacterium Tuberculosis.” *The Journal of Antimicrobial Chemotherapy* 35 (2): 345–48. https://doi.org/10.1093/jac/35.2.345.

Bostanabad, Saeed Zaker, Ahmadreza Bahrmand, Leonid P. Titov, and Mohammad Taghikhani. 2007. “Identification of Mutations in the rpoB Encoding the RNA Polymerase Beta Subunit in Rifampicine-Resistant Mycobacterium Tuberculosis Strains from Iran.” *Tuberkuloz Ve Toraks* 55 (4): 370–77.

Campbell, Patricia J., Glenn P. Morlock, R. David Sikes, Tracy L. Dalton, Beverly Metchock, Angela M. Starks, Delaina P. Hooks, Lauren S. Cowan, Bonnie B. Plikaytis, and James E. Posey. 2011. “Molecular Detection of Mutations Associated with First- and Second-Line Drug Resistance Compared with Conventional Drug Susceptibility Testing of Mycobacterium Tuberculosis.” *Antimicrobial Agents and Chemotherapy* 55 (5): 2032–41. https://doi.org/10.1128/AAC.01550-10.

Cavusoglu, Cengiz, Suleyha Hilmioglu, Sevinc Guneri, and Altinay Bilgic. 2002. “Characterization of rpoB Mutations in Rifampin-Resistant Clinical Isolates of Mycobacterium Tuberculosis from Turkey by DNA Sequencing and Line Probe Assay.” *Journal of Clinical Microbiology* 40 (12): 4435–38. https://doi.org/10.1128/JCM.40.12.4435-4438.2002.

ElMaraachli, W., M. Slater, Z. L. Berrada, S.-Y. G. Lin, A. Catanzaro, E. Desmond, C. Rodrigues, et al. 2015. “Predicting Differential Rifamycin Resistance in Clinical Mycobacterium Tuberculosis Isolates by Specific rpoB Mutations.” *The International Journal of Tuberculosis and Lung Disease: The Official Journal of the International Union Against Tuberculosis and Lung Disease* 19 (10): 1222–26. https://doi.org/10.5588/ijtld.14.0936.

Heep, M., U. Rieger, D. Beck, and N. Lehn. 2000. “Mutations in the Beginning of the rpoB Gene Can Induce Resistance to Rifamycins in Both Helicobacter Pylori and Mycobacterium Tuberculosis.” *Antimicrobial Agents and Chemotherapy* 44 (4): 1075–77. https://doi.org/10.1128/AAC.44.4.1075-1077.2000.

Heym, B., N. Honoré, C. Truffot-Pernot, A. Banerjee, C. Schurra, W. R. Jacobs, J. D. van Embden, J. H. Grosset, and S. T. Cole. 1994. “Implications of Multidrug Resistance for the Future of Short-Course Chemotherapy of Tuberculosis: A Molecular Study.” *Lancet (London, England)* 344 (8918): 293–98. https://doi.org/10.1016/s0140-6736(94)91338-2.

Htike Min, Pyar Kyi, Pannamthip Pitaksajjakul, Natthakan Tipkrua, Waranya Wongwit, Pornrutsami Jintaridh, and Pongrama Ramasoota. 2014. “Novel Mutation Detection IN rpoB OF Rifampicin-Resistant Mycobacterium Tuberculosis Using Pyrosequencing.” *The Southeast Asian Journal of Tropical Medicine and Public Health* 45 (4): 843–52.

Jamieson, F. B., J. L. Guthrie, A. Neemuchwala, O. Lastovetska, R. G. Melano, and C. Mehaffy. 2014. “Profiling of rpoB Mutations and MICs for Rifampin and Rifabutin in Mycobacterium Tuberculosis.” *Journal of Clinical Microbiology* 52 (6): 2157–62. https://doi.org/10.1128/JCM.00691-14.

Jing, Wei, Yu Pang, Zhaojing Zong, Jing Wang, Ru Guo, Fengmin Huo, Guanglu Jiang, Yifeng Ma, Hairong Huang, and Naihui Chu. 2017. “Rifabutin Resistance Associated with Double Mutations in rpoB Gene in Mycobacterium Tuberculosis Isolates.” *Frontiers in Microbiology* 8: 1768. https://doi.org/10.3389/fmicb.2017.01768.

Kambli, Priti, Kanchan Ajbani, Meeta Sadani, Chaitali Nikam, Anjali Shetty, Zarir Udwadia, Sophia B. Georghiou, Timothy C. Rodwell, Antonino Catanzaro, and Camilla Rodrigues. 2015. “Defining Multidrug-Resistant Tuberculosis: Correlating GenoType MTBDRplus Assay Results with Minimum Inhibitory Concentrations.” *Diagnostic Microbiology and Infectious Disease* 82 (1): 49–53. https://doi.org/10.1016/j.diagmicrobio.2015.01.009.

Kim, B. J., S. Y. Kim, B. H. Park, M. A. Lyu, I. K. Park, G. H. Bai, S. J. Kim, C. Y. Cha, and Y. H. Kook. 1997. “Mutations in the rpoB Gene of Mycobacterium Tuberculosis That Interfere with PCR-Single-Strand Conformation Polymorphism Analysis for Rifampin Susceptibility Testing.” *Journal of Clinical Microbiology* 35 (2): 492–94. https://doi.org/10.1128/jcm.35.2.492-494.1997.

Madania, Ammar, Maya Habous, Hana Zarzour, Ifad Ghoury, and Barea Hebbo. 2012. “Characterization of Mutations Causing Rifampicin and Isoniazid Resistance of Mycobacterium Tuberculosis in Syria.” *Polish Journal of Microbiology* 61 (1): 23–32.

Matsui, Tania, Juliana Maíra Watanabe Pinhata, Michelle Christiane da Silva Rabello, Angela Pires Brandão, Lucilaine Ferrazoli, Sylvia Cardoso Leão, Cristina Viana-Niero, and Rosangela Siqueira de Oliveira. 2020. “Frequency of First and Second-Line Drug Resistance-Associated Mutations among Resistant Mycobacterium Tuberculosis Clinical Isolates from São Paulo, Brazil.” *Memorias Do Instituto Oswaldo Cruz* 115: e200055. https://doi.org/10.1590/0074-02760200055.

Miotto, Paolo, Andrea M. Cabibbe, Emanuele Borroni, Massimo Degano, and Daniela M. Cirillo. 2018. “Role of Disputed Mutations in the rpoB Gene in Interpretation of Automated Liquid MGIT Culture Results for Rifampin Susceptibility Testing of Mycobacterium Tuberculosis.” *Journal of Clinical Microbiology* 56 (5): e01599-17. https://doi.org/10.1128/JCM.01599-17.

Mvelase, Nomonde R., Melendhran Pillay, Wilbert Sibanda, Jacqueline N. Ngozo, James C. M. Brust, and Koleka P. Mlisana. 2019. “rpoB Mutations Causing Discordant Rifampicin Susceptibility in Mycobacterium Tuberculosis: Retrospective Analysis of Prevalence, Phenotypic, Genotypic, and Treatment Outcomes.” *Open Forum Infectious Diseases* 6 (4): ofz065. https://doi.org/10.1093/ofid/ofz065.

Nosova, Elena Y., Danila V. Zimenkov, Anastasia A. Khakhalina, Alexandra I. Isakova, Ludmila Y. Krylova, Marina V. Makarova, Ksenia Y. Galkina, et al. 2016. “A Comparison of the Sensititre MycoTB Plate, the Bactec MGIT 960, and a Microarray-Based Molecular Assay for the Detection of Drug Resistance in Clinical Mycobacterium Tuberculosis Isolates in Moscow, Russia.” *PloS One* 11 (11): e0167093. https://doi.org/10.1371/journal.pone.0167093.

Otchere, I. D., A. Asante-Poku, S. Osei-Wusu, A. Baddoo, E. Sarpong, A. H. Ganiyu, S. Y. Aboagye, et al. 2016. “Detection and Characterization of Drug-Resistant Conferring Genes in Mycobacterium Tuberculosis Complex Strains: A Prospective Study in Two Distant Regions of Ghana.” *Tuberculosis (Edinburgh, Scotland)* 99 (July): 147–54. https://doi.org/10.1016/j.tube.2016.05.014.

Rodwell, Timothy C., Faramarz Valafar, James Douglas, Lishi Qian, Richard S. Garfein, Ashu Chawla, Jessica Torres, et al. 2014. “Predicting Extensively Drug-Resistant Mycobacterium Tuberculosis Phenotypes with Genetic Mutations.” *Journal of Clinical Microbiology* 52 (3): 781–89. https://doi.org/10.1128/JCM.02701-13.

Rukasha, Ivy, Halima M. Said, Shaheed V. Omar, Hendrik Koornhof, Andries W. Dreyer, Alfred Musekiwa, Harry Moultrie, et al. 2016. “Correlation of rpoB Mutations with Minimal Inhibitory Concentration of Rifampin and Rifabutin in Mycobacterium Tuberculosis in an HIV/AIDS Endemic Setting, South Africa.” *Frontiers in Microbiology* 7: 1947. https://doi.org/10.3389/fmicb.2016.01947.

Sandgren, Andreas, Michael Strong, Preetika Muthukrishnan, Brian K. Weiner, George M. Church, and Megan B. Murray. 2009. “Tuberculosis Drug Resistance Mutation Database.” *PLoS Medicine* 6 (2): e2. https://doi.org/10.1371/journal.pmed.1000002.

Schilke, K., K. Weyer, G. Bretzel, B. Amthor, J. Brandt, V. Sticht-Groh, P. B. Fourie, and W. H. Haas. 1999. “Universal Pattern of RpoB Gene Mutations among Multidrug-Resistant Isolates of Mycobacterium Tuberculosis Complex from Africa.” *The International Journal of Tuberculosis and Lung Disease: The Official Journal of the International Union Against Tuberculosis and Lung Disease* 3 (7): 620–26.

Singh, Binit Kumar, Rohini Sharma, Parul Kodan, Manish Soneja, Pankaj Jorwal, Neeraj Nischal, Ashutosh Biswas, Sanjay Sarin, Ranjani Ramachandran, and Naveet Wig. 2020. “Diagnostic Evaluation of Non-Interpretable Results Associated with rpoB Gene in Genotype MTBDRplus Ver 2.0.” *Tuberculosis and Respiratory Diseases* 83 (4): 289–94. https://doi.org/10.4046/trd.2020.0039.

Sirgel, Frederick A., Robin M. Warren, Erik C. Böttger, Marisa Klopper, Thomas C. Victor, and Paul D. van Helden. 2013. “The Rationale for Using Rifabutin in the Treatment of MDR and XDR Tuberculosis Outbreaks.” *PloS One* 8 (3): e59414. https://doi.org/10.1371/journal.pone.0059414.

Solari, L., D. Santos-Lazaro, and Z. M. Puyen. 2020. “Mutations in Mycobacterium Tuberculosis Isolates with Discordant Results for Drug-Susceptibility Testing in Peru.” *International Journal of Microbiology* 2020: 8253546. https://doi.org/10.1155/2020/8253546.

Somoskovi, Akos, Vanessa Deggim, Diana Ciardo, and Guido V. Bloemberg. 2013. “Diagnostic Implications of Inconsistent Results Obtained with the Xpert MTB/Rif Assay in Detection of Mycobacterium Tuberculosis Isolates with an rpoB Mutation Associated with Low-Level Rifampin Resistance.” *Journal of Clinical Microbiology* 51 (9): 3127–29. https://doi.org/10.1128/JCM.01377-13.

Springer, Burkhard, Katja Lucke, Romana Calligaris-Maibach, Claudia Ritter, and Erik C. Böttger. 2009. “Quantitative Drug Susceptibility Testing of Mycobacterium Tuberculosis by Use of MGIT 960 and EpiCenter Instrumentation.” *Journal of Clinical Microbiology* 47 (6): 1773–80. https://doi.org/10.1128/JCM.02501-08.

Stavrum, Ruth, Vithal Prasad Myneedu, Virendra K. Arora, Niyaz Ahmed, and Harleen M. S. Grewal. 2009. “In-Depth Molecular Characterization of Mycobacterium Tuberculosis from New Delhi--Predominance of Drug Resistant Isolates of the ‘modern’ (TbD1) Type.” *PloS One* 4 (2): e4540. https://doi.org/10.1371/journal.pone.0004540.

Suthum, Krairerk, Worada Samosornsuk, and Seksun Samosornsuk. 2020. “Characterization of katG, inhA, rpoB and pncA in Mycobacterium Tuberculosis Isolates from MDR-TB Risk Patients in Thailand.” *Journal of Infection in Developing Countries* 14 (3): 268–76. https://doi.org/10.3855/jidc.11974.

Taniguchi, H., H. Aramaki, Y. Nikaido, Y. Mizuguchi, M. Nakamura, T. Koga, and S. Yoshida. 1996. “Rifampicin Resistance and Mutation of the rpoB Gene in Mycobacterium Tuberculosis.” *FEMS Microbiology Letters* 144 (1): 103–8. https://doi.org/10.1111/j.1574-6968.1996.tb08515.x.

Telenti, A., P. Imboden, F. Marchesi, D. Lowrie, S. Cole, M. J. Colston, L. Matter, K. Schopfer, and T. Bodmer. 1993. “Detection of Rifampicin-Resistance Mutations in Mycobacterium Tuberculosis.” *Lancet (London, England)* 341 (8846): 647–50. https://doi.org/10.1016/0140-6736(93)90417-f.

Torrea, Gabriela, Kamela C. S. Ng, Armand Van Deun, Emmanuel André, Justine Kaisergruber, Willy Ssengooba, Christel Desmaretz, et al. 2019. “Variable Ability of Rapid Tests to Detect Mycobacterium Tuberculosis rpoB Mutations Conferring Phenotypically Occult Rifampicin Resistance.” *Scientific Reports* 9 (1): 11826. https://doi.org/10.1038/s41598-019-48401-z.

Van Deun, Armand, Kya J. M. Aung, Valentin Bola, Rossin Lebeke, Mohamed Anwar Hossain, Willem Bram de Rijk, Leen Rigouts, Aysel Gumusboga, Gabriela Torrea, and Bouke C. de Jong. 2013. “Rifampin Drug Resistance Tests for Tuberculosis: Challenging the Gold Standard.” *Journal of Clinical Microbiology* 51 (8): 2633–40. https://doi.org/10.1128/JCM.00553-13.

Vargas, Ana Paula, Angela A. Rios, Louis Grandjean, Daniela E. Kirwan, Robert H. Gilman, Patricia Sheen, and Mirko J. Zimic. 2020. “Determination of Potentially Novel Compensatory Mutations in Rpoc Associated with Rifampin Resistance and Rpob Mutations in Mycobacterium Tuberculosis Clinical Isolates from Peru.” *International Journal of Mycobacteriology* 9 (2): 121–37. https://doi.org/10.4103/ijmy.ijmy_27_20.

Walker, Timothy M., Paolo Miotto, Claudio U. Köser, Philip W. Fowler, Jeff Knaggs, Zamin Iqbal, Martin Hunt, et al. 2022. “The 2021 WHO Catalogue of Mycobacterium Tuberculosis Complex Mutations Associated with Drug Resistance: A Genotypic Analysis.” *The Lancet. Microbe* 3 (4): e265–73. https://doi.org/10.1016/S2666-5247(21)00301-3.

Wang, Qingzhong, Jun Yue, Lu Zhang, Ying Xu, Jiazhen Chen, Min Zhang, Bingdong Zhu, Hongyan Wang, and Honghai Wang. 2007. “A Newly Identified 191A/C Mutation in the Rv2629 Gene That Was Significantly Associated with Rifampin Resistance in Mycobacterium Tuberculosis.” *Journal of Proteome Research* 6 (12): 4564–71. https://doi.org/10.1021/pr070242z.

Williamson, D. A., S. A. Roberts, J. E. Bower, R. Vaughan, S. Newton, O. Lowe, C. A. Lewis, and J. T. Freeman. 2012. “Clinical Failures Associated with rpoB Mutations in Phenotypically Occult Multidrug-Resistant Mycobacterium Tuberculosis.” *The International Journal of Tuberculosis and Lung Disease: The Official Journal of the International Union Against Tuberculosis and Lung Disease* 16 (2): 216–20. https://doi.org/10.5588/ijtld.11.0178.

Yang, B., H. Koga, H. Ohno, K. Ogawa, M. Fukuda, Y. Hirakata, S. Maesaki, K. Tomono, T. Tashiro, and S. Kohno. 1998. “Relationship between Antimycobacterial Activities of Rifampicin, Rifabutin and KRM-1648 and rpoB Mutations of Mycobacterium Tuberculosis.” *The Journal of Antimicrobial Chemotherapy* 42 (5): 621–28. https://doi.org/10.1093/jac/42.5.621.

Zaczek, Anna, Anna Brzostek, Ewa Augustynowicz-Kopec, Zofia Zwolska, and Jaroslaw Dziadek. 2009. “Genetic Evaluation of Relationship between Mutations in rpoB and Resistance of Mycobacterium Tuberculosis to Rifampin.” *BMC Microbiology* 9 (January): 10. https://doi.org/10.1186/1471-2180-9-10.

Zignol, Matteo, Andrea Maurizio Cabibbe, Anna S. Dean, Philippe Glaziou, Natavan Alikhanova, Cecilia Ama, Sönke Andres, et al. 2018. “Genetic Sequencing for Surveillance of Drug Resistance in Tuberculosis in Highly Endemic Countries: A Multi-Country Population-Based Surveillance Study.” *The Lancet. Infectious Diseases* 18 (6): 675–83. https://doi.org/10.1016/S1473-3099(18)30073-2.

### Table S2.

Statistically significant list of genes based on dN/dS, d2N/d1N, and d2N/dS ratios statistics. Association of measures with drug resistance (a), gene-gene pairwise comparisons for source mutation frequencies (b), gene-gene pairwise comparisons for phylogenetically-adjusted frequencies (c), and individual codon mutation frequencies deviations from the expected values (d) approaches were used.

| Gene | Resistance, or resistance-associated gene* | Number of S mutations | Number of 1N mutations | Number of 2N mutations | Odds ratio | | | -*log*(*p*-value) * | | |
| --- | --- | --- | --- | --- | --- | --- | --- | --- | --- | --- |
|  |  |  |  |  | dN/ dS | d2N/ d1N | d2N/ dS | dN/ dS | d2N/ d1N | d2N/ dS |
| *pncA* | R | 25 | 886 | 0 | 17.40 | 0.00 | 0.00 | **71.30** | **4.30** |  |
| *rpoB* | R | 108 | 2108 | 43 | 10.44 | 3.76 | 38.67 | **151.73** |  | **28.16** |
| *cut1* | R | 9 | 160 | 0 | 9.29 | 0.00 | 0.00 | **11.07** | **8.67** |  |
| *pstP* |  | 89 | 1705 | 0 | 8.67 | 0.00 | 0.00 | **99.79** | **6.81** |  |
| *dipZ* |  | 83 | 2599 | 1 | 7.78 | 0.01 | 0.12 | **90.17** | **17.27** |  |
| *cyp138* |  | 64 | 108 | 350 | 6.68 | 19.20 | 35.34 | **37.43** | **62.27** | **84.25** |
| *katG* | R | 100 | 1396 | 23 | 5.79 | 2.21 | 12.65 | **61.19** |  | **9.87** |
| *Rv0988* |  | 211 | 271 | 244 | 1.81 | 10.14 | 10.51 | **6.99** | **41.54** | **40.39** |
| *Rv2082* |  | 819 | 1223 | 0 | 1.25 | 0.00 | 0.00 | 3.40 | **8.45** | **6.86** |
| *Rv2024c* |  | 102 | 189 | 42 | 0.94 | 52.00 | 40.35 | 0.10 | **12.30** | **9.23** |
| *recC* |  | 807 | 183 | 1 | 0.13 | 0.08 | 0.01 | **92.24** |  | **17.63** |
| *Rv1190* |  | 347 | 50 | 0 | 0.10 | 0.00 | 0.00 | **42.76** |  | **5.30** |
| *mce1F* |  | 2201 | 205 | 1 | 0.06 | 0.23 | 0.01 | **200.00** |  | **14.88** |
| *Rv1977* |  | 684 | 68 | 0 | 0.05 | 0.00 | 0.00 | **128.47** |  | **5.68** |
| *hsaB* |  | 24 | 1553 | 0 | 36.73 | 0.00 | 0.00 | **161.74** |  |  |
| *Rv0192* |  | 42 | 1520 | 1 | 20.45 | 0.11 | 2.24 | **140.61** |  |  |
| *cobD* |  | 50 | 2043 | 0 | 12.76 | 0.00 | 0.00 | **105.15** |  |  |
| *murD* |  | 55 | 1171 | 1 | 12.13 | 0.13 | 1.62 | **93.14** |  |  |
| *Rv0395* |  | 16 | 326 | 0 | 12.04 |  |  | **26.84** |  |  |
| *rpsL* | R | 26 | 572 | 2 | 8.73 | 0.37 | 3.27 | **30.87** |  |  |
| *Rv0315* |  | 29 | 361 | 0 | 7.95 | 0.00 | 0.00 | **26.09** | 0.80 | 0.00 |
| *Rv2668* |  | 12 | 222 | 0 | 7.90 |  |  | **12.26** | 0.00 | 0.00 |
| *rbsK* |  | 28 | 371 | 0 | 7.32 | 0.00 | 0.00 | **23.32** |  |  |
| *Rv1760* |  | 33 | 521 | 0 | 7.29 | 0.00 | 0.00 | **28.69** |  |  |
| *proB* |  | 53 | 871 | 0 | 7.07 | 0.00 | 0.00 | **44.49** |  |  |
| *Rv0538* |  | 58 | 680 | 0 | 6.92 | 0.00 | 0.00 | **42.86** |  |  |
| *Rv0064* |  | 149 | 1842 | 1 | 6.24 | 0.10 | 0.64 | **97.02** |  |  |
| *Rv0318c* |  | 31 | 342 | 1 | 6.11 | 0.10 | 0.61 | **19.39** | 2.00 | 0.00 |
| *Rv3657c* |  | 17 | 105 | 0 | 5.98 |  |  | **8.45** | 0.00 | 0.00 |
| *Rv0398c* |  | 23 | 437 | 0 | 5.69 |  |  | **15.08** | 0.00 | 0.00 |
| *bioD* |  | 27 | 241 | 0 | 5.06 |  |  | **12.31** | 0.00 | 0.00 |
| *ponA1* |  | 88 | 811 | 0 | 5.02 | 0.00 | 0.00 | **38.92** |  |  |
| *Rv1230c* |  | 39 | 253 | 1 | 4.91 | 0.66 | 3.23 | **15.07** | 0.00 | 0.37 |
| *gid* | R | 79 | 682 | 7 | 4.54 | 0.64 | 2.91 | **29.68** |  |  |
| *phoR* |  | 49 | 376 | 1 | 4.27 | 0.24 | 1.03 | **16.28** | 0.50 | 0.00 |
| *murG* |  | 53 | 367 | 0 | 4.09 | 0.00 | 0.00 | **15.90** |  |  |
| *Rv0218* |  | 42 | 424 | 0 | 3.96 | 0.00 | 0.00 | **13.62** | 0.71 | 0.00 |
| *ddlA* |  | 37 | 194 | 0 | 3.86 |  |  | **9.30** | 0.00 | 0.00 |
| *ethA* | R | 64 | 447 | 3 | 3.44 | 0.32 | 1.13 | **14.40** | 0.95 | 0.00 |
| *Rv1129c* | R | 39 | 212 | 0 | 3.42 | 0.00 | 0.00 | **8.15** | 0.75 | 0.00 |
| *gyrA* | R | 194 | 1308 | 2 | 3.34 | 0.19 | 0.64 | **39.66** |  |  |
| *whiB6* | R | 33 | 230 | 1 | 3.27 | 0.26 | 0.85 | **6.95** | 0.49 | 0.00 |
| *Rv2264c* |  | 75 | 423 | 0 | 3.14 | 0.00 | 0.00 | **13.22** | 0.37 | 0.00 |
| *rpoC* | R | 136 | 786 | 3 | 3.04 | 0.29 | 0.88 | **22.32** |  |  |
| *cycA* |  | 80 | 239 | 0 | 2.72 | 0.00 | 0.00 | **8.38** | 0.77 | 0.26 |
| *embB* | R | 154 | 1485 | 0 | 2.70 | 0.00 | 0.00 | **22.48** | 1.00 | 0.25 |
| *Rv1461* |  | 80 | 349 | 0 | 2.47 | 0.00 | 0.00 | **7.92** | 0.36 | 0.00 |
| *Rv3433c* |  | 993 | 1010 | 0 | 0.65 | 0.00 | 0.00 | **10.45** | 1.74 | 2.48 |
| *aftD* |  | 916 | 825 | 1 | 0.56 | 0.11 | 0.06 | **16.04** | 2.03 | 3.56 |
| *ppsD* |  | 261 | 217 | 0 | 0.47 | 0.00 | 0.00 | **7.84** | 0.29 | 0.79 |
| *mmpL4* |  | 151 | 125 | 0 | 0.37 | 0.00 | 0.00 | **7.50** | 0.00 | 0.44 |
| *recD* |  | 196 | 107 | 0 | 0.35 | 0.00 | 0.00 | **9.51** | 0.27 | 0.84 |
| *dctA* |  | 342 | 83 | 0 | 0.33 | 0.00 | 0.00 | **11.96** | 0.00 | 0.38 |
| *serA1* |  | 120 | 73 | 0 | 0.29 | 0.00 | 0.00 | **8.05** | 0.79 | 2.25 |
| *hemE* |  | 145 | 41 | 0 | 0.29 |  |  | **7.32** | 0.00 | 0.00 |
| *Rv2081c* |  | 91 | 54 | 2 | 0.27 | 1.85 | 0.49 | **6.95** | 0.21 | 0.22 |
| *eccC5* |  | 193 | 93 | 0 | 0.26 | 0.00 | 0.00 | **13.79** | 0.26 | 0.92 |
| *Rv1501* |  | 149 | 74 | 0 | 0.25 |  |  | **11.82** | 0.00 | 0.00 |
| *Rv2319c* |  | 160 | 56 | 1 | 0.22 | 0.78 | 0.17 | **12.97** | 0.00 | 0.91 |
| *Rv3528c* |  | 374 | 92 | 0 | 0.18 | 0.00 | 0.00 | **33.61** |  |  |
| *Rv0575c* |  | 218 | 81 | 1 | 0.17 | 1.27 | 0.21 | **24.08** | 0.00 | 0.66 |
| *Rv0376c* |  | 2498 | 336 | 1 | 0.17 | 1.25 | 0.21 | **170.00** |  |  |
| *Rv0986* |  | 218 | 60 | 1 | 0.17 | 2.90 | 0.48 | **22.13** | 0.35 | 0.26 |
| *ruvB* |  | 176 | 41 | 0 | 0.14 | 0.00 | 0.00 | **19.36** | 0.00 | 0.98 |
| *Rv2052c* |  | 852 | 131 | 0 | 0.10 | 0.00 | 0.00 | **113.93** |  |  |
| *Rv3228* | R | 521 | 75 | 0 | 0.09 | 0.00 | 0.00 | **68.81** |  |  |
| *Rv1319c* |  | 162 | 23 | 7 | 0.06 | 21.46 | 1.06 | **31.84** |  |  |
| *Rv1762c* |  | 144 | 26 | 0 | 0.04 | 0.00 | 0.00 | **34.81** |  |  |
| *relG* |  | 1339 | 12 | 0 | 0.01 | 0.00 | 0.00 | **181.38** |  |  |
| *Rv1573* |  | 667 | 8 | 0 | 0.01 | 0.00 | 0.00 | **142.67** |  |  |

### Table S3.

Statistically significant list of genes of *Neisseria gonorrhoeae* based on dN/dS, d2N/d1N, and d2N/dS ratios statistics.

| Gene | Number of S mutations | Number of 1N mutations | Number of 2N mutations | Odds ratio | | | -*log*(*p*-value) * | | |
| --- | --- | --- | --- | --- | --- | --- | --- | --- | --- |
|  |  |  |  | dN/ dS | d2N/ d1N | d2N/ dS | dN/ dS | d2N/ d1N | d2N/ dS |
| NGK_RS00205 | 363 | 88 | 23 | 4.4 | 14.9 | 62.2 | **24.3** | 3.8 | **11.0** |
| NGK_RS06565 | 638 | 78 | 24 | 7.1 | 5.0 | 34.7 | **52.8** | 2.4 | **13.1** |
| NGK_RS06660 | 538 | 135 | 18 | 3.7 | 6.0 | 21.9 | **28.0** | 2.3 | **7.3** |
| NGK_RS06730 | 2255 | 381 | 35 | 3.9 | 6.3 | 24.2 | **97.2** | **4.6** | **15.3** |
| NGK_RS08025 | 2766 | 295 | 28 | 6.4 | 1.9 | 11.8 | **180.9** | 1.0 | **11.9** |
| NGK_RS08860 | 1133 | 288 | 20 | 4.1 | 1.8 | 7.3 | **67.9** | 0.6 | **5.8** |
| NGK_RS09060 | 886 | 324 | 23 | 2.7 | 5.4 | 14.4 | **31.9** | 2.7 | **7.5** |
| NGK_RS09405 | 329 | 50 | 24 | 10.2 | 11.9 | 114.2 | **45.3** | 2.9 | **15.1** |
| NGK_RS09735 | 2564 | 178 | 13 | 8.9 | 1.2 | 11.0 | **189.5** | 0.1 | **6.3** |
| NGK_RS09870 | 1874 | 266 | 11 | 4.7 | 2.5 | 11.8 | **95.9** | 0.8 | **4.6** |
| NGK_RS11680 | 613 | 169 | 23 | 4.2 | 7.1 | 28.9 | **39.8** | 2.9 | **9.9** |
| NGK_RS14155 | 117 | 25 | 12 | 5.9 | 7.3 | 39.4 | **11.1** | 1.5 | **5.7** |
| NGK_RS15570 | 2110 | 206 | 27 | 6.3 | 1.9 | 11.6 | **128.5** | 0.9 | **11.4** |
| NGK_RS15575 | 2001 | 116 | 29 | 10.0 | 2.8 | 27.8 | **148.5** | 2.0 | **18.0** |
| NGK_RS15580 | 1692 | 127 | 10 | 8.8 | 1.6 | 14.3 | **129.2** | 0.2 | **5.4** |
